# CLCNet: a contrastive learning and chromosome-aware network for genomic prediction in plants

**DOI:** 10.1101/2024.12.29.630569

**Authors:** Jiangwei Huang, Zhihan Yang, Mou Yin, Chao Li, Jinmin Li, Yu Wang, Lu Huang, Miaomiao Li, Chengzhi Liang, Fei He, Rongcheng Han, Yuqiang Jiang

## Abstract

Genomic selection (GS) leverages genome-wide markers and phenotypes to predict breeding values, with its effectiveness largely dependent on the accuracy of genomic prediction (GP) models. However, GP methods often struggle to capture inter-individual variability and are limited by the curse of dimensionality, where the number of SNPs far exceeds the sample size. To address these challenges, we present CLCNet (Contrastive Learning and Chromosome-aware Network), a novel deep learning framework that integrates contrastive learning and chromosome-aware feature modeling. CLCNet comprises two key components: (i) a contrastive learning module that enhances the model’s ability to capture fine-grained, genotype-dependent phenotypic differences among individuals, and (ii) a chromosome-aware module that captures structured feature selection at both chromosome and genome levels, thereby distilling the most informative SNPs. We evaluated CLCNet across four crop species, covering ten agronomically important traits, and compared it with a diverse set of classical linear, machine learning, and deep learning models. CLCNet achieved superior prediction performance, with statistically significant improvements in Pearson correlation coefficient (PCC), ranging from 0.34% to 12.19% over baseline, together with reduced mean squared error (MSE). Performance gains were more pronounced for traits with moderate linkage disequilibrium (LD; *r*^2^ = 0.21-0.36) and high heritability (*h*^2^ > 0.66), such as those in maize, rapeseed, and soybean. For cotton traits characterized by high LD (*r*^2^ = 0.74) and lower heritability (*h*^2^ < 0.50), CLCNet maintained robust performance without degradation. Overall, these results demonstrate that CLCNet is an effective framework for improving genomic prediction accuracy and holds strong potential for practical applications in plant breeding.

**Short abstract:** CLCNet is a novel deep learning framework for genomic prediction that integrates contrastive learning with chromosome-aware feature selection. By jointly modeling inter-individual genotype–phenotype variation and chromosomal genomic structure, CLCNet improves prediction accuracy under high-dimensional, low-sample-size conditions. Across four crop species and ten agronomic traits, CLCNet consistently outperformed classical statistical, machine learning, and existing deep learning models. The framework also identified biologically relevant SNPs and candidate genes, demonstrating its potential for practical applications in genomic selection and computational plant breeding.

**Key points:**

- We propose CLCNet, a multi-task deep learning framework that integrates contrastive learning with chromosome-aware feature selection for genomic prediction, under high-dimensional, low-sample-size conditions.
- The chromosome-aware module explicitly exploits genomic structural information to select representative and informative SNPs across chromosomes.
- Contrastive learning improves model robustness by stabilizing representation learning and reducing the influence of random effects across samples.
- By complementing GWAS analyses, CLCNet provides additional insights into genotype–phenotype relationships with potential relevance for gene discovery.

**Biographical Note:** **Jiangwei Huang** is a PhD candidate at the Institute of Genetics and Developmental Biology, Chinese Academy of Sciences. His research focuses on genomic prediction, deep learning, and computational plant breeding.

**Zhihan Yang** is a PhD candidate at the Institute of Genetics and Developmental Biology, Chinese Academy of Sciences. Her research interests include genomic prediction and bioinformatics.

**Rongcheng Han** is an associate professor at the Institute of Genetics and Developmental Biology, Chinese Academy of Sciences. His research focuses on bioinformatics and plant phenomics.

**Yuqiang Jiang** is a professor at the Institute of Genetics and Developmental Biology, Chinese Academy of Sciences. His research interests include plant genomics, plant phenomics and genetic improvement.

**Organization description:** The Institute of Genetics and Developmental Biology, Chinese Academy of Sciences, is a leading research institute focusing on genetics, genomics, molecular breeding, bioinformatics, and systems biology in plants and animals.

## Introduction

Genomic selection (GS) has become an increasingly important approach in modern plant breeding by enabling the prediction of genomic estimated breeding values (GEBVs) through the integration of genome-wide molecular markers with phenotypic data [1]. Unlike marker-assisted selection (MAS), which primarily focuses on loci with large effects, GS captures the collective contribution of a large number of loci with small to moderate effects distributed across the genome, thereby providing a more comprehensive and accurate framework for the prediction of agronomic traits [2–4]. A wide range of genomic prediction (GP) models has been developed, including classical linear models (e.g., rrBLUP and GBLUP), and Bayesian regression approaches (e.g., Bayesian A/B/C/Cπ), as well as machine learning methods such as support vector regression and random forest regression [5–11].

More recently, deep learning (DL) has garnered increasing attention for its strong capacity to model nonlinear genotype-phenotype relationships through hierarchical architectures [12–14]. Existing DL-based GP models span diverse paradigms, including single-task, multi-modal, and transfer learning [15, 16]. Representative single-task learning models include DeepGS [17], DLGWAS [18], SoyDNGP [19], DNNGP [20], and Cropformer [21], while DEM [22], DeepCCR [23], and GEFormer [24] exemplify multi-modal learning, and TrG2P [25] represents transfer learning model-based GP. However, multi-task learning, despite its potential to jointly model multiple traits and enable information sharing, remains underexplored in GP. In addition, most DL-based approaches do not adequately capture fine-grained inter-individual variation in genotype–phenotype relationships.

Moreover, GP performance remains constrained by the “curse of dimensionality”, where the number of SNPs far exceeds the number of samples. This considerable imbalance (e.g., >100,000 features vs. <10,000 samples) frequently leads to overfitting, instability, and reduced generalizability of predictive models, particularly in the presence of genomic noise or limited phenotype data [10, 26, 27]. Consequently, effective feature selection has been recognized as a critical step. However, most methods treat SNPs as independent features and overlook the inherent chromosomal structure of the genome, thereby failing to capture both local and long-range genetic interactions, which plays a crucial role across the genome [28–30].

To address these challenges, we propose CLCNet (Contrastive Learning and Chromosome-aware Network), a multi-task deep learning framework that integrates two key components: (i) a contrastive learning module to capture inter-individual differences in genotype–phenotype relationships, and (ii) a chromosome-aware module that performs hierarchical SNP selection at both intra- and inter-chromosomal levels. We evaluated CLCNet across four crop species-maize (*Zea mays*), cotton (*Gossypium hirsutum*), rapeseed (*Brassica napus*), and soybean (*Glycine max*), across ten distinct traits, and compared it with three classical linear models (rrBLUP, Bayesian Ridge, Bayesian Lasso), two machine learning models (LightGBM, SVR), and two deep learning models (DNNGP, DeepGS). Our results collectively demonstrate that CLCNet provides a robust and biologically informed framework for improving GP and shows potential for accelerating genetic gain in plant breeding.

## Results

### Architectural design of CLCNet

CLCNet is a multi-task learning framework designed to predict phenotypic traits from high-dimensional SNP data while jointly learning low-dimensional SNP embeddings as an auxiliary task. The core architecture adopts a dual-pathway backbone, consisting of a non-linear pathway with three fully connected layers and a parallel linear pathway. Their outputs are combined through a residual connection, enabling the model to preserve linear additive effects while capturing non-linear interactions (Fig. 1 A). The shared backbone representations are then fed into two task-specific branches: a regression head for phenotypic prediction, trained using mean squared error (MSE), and an auxiliary head that generates SNP embeddings for contrastive learning.

**Fig. 1.**
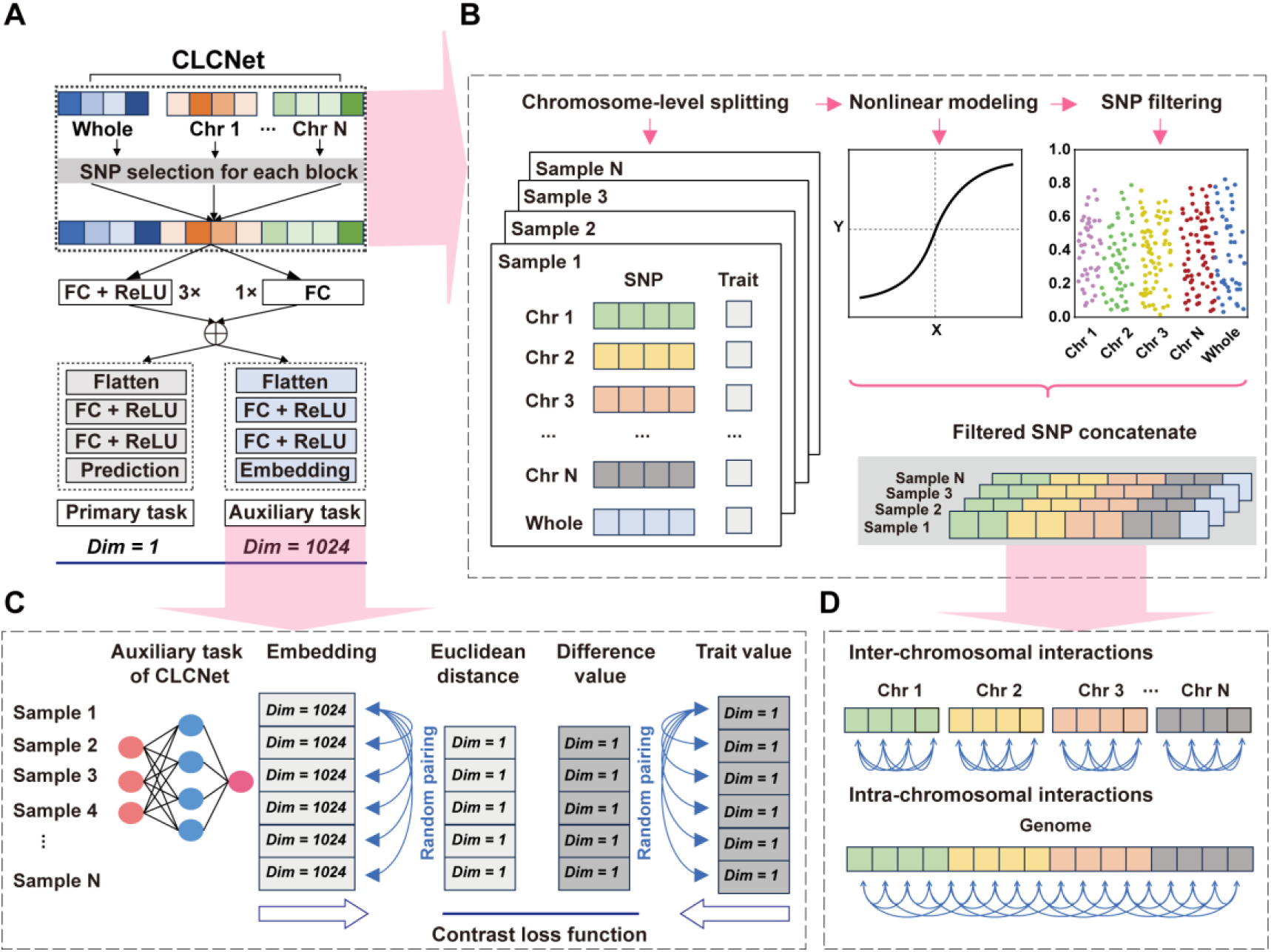
The Framework of CLCNet Model. **(A) CLCNet architecture.** FC denotes fully connected layers, and ReLU denotes the activation function. “3×” and “1×” indicate the number of repeated blocks. “Dim” refers to feature dimensionality; the primary task (trait prediction) outputs a single value, whereas the auxiliary task (embedding) outputs a 1024-dimensional vector. **(B) Chromosome-aware module.** Genome-wide SNPs are first partitioned into chromosome-specific groups while retaining the original genome-wide SNP set. Nonlinear feature selection methods, such as LightGBM, are then applied to identify SNPs with positive contributions. The selected local and global SNPs were merged into a unified feature set, and redundant SNPs were removed to construct the final input features for the model. **(C) Contrastive learning module.** For each sample, the 1024-dimensional embedding generated by the auxiliary task is used for contrastive learning. Randomly paired samples are compared by computing the Euclidean distance between embeddings, while the absolute difference between their phenotypic values serves as a one-dimensional reference. Both quantities are incorporated into a customized contrastive loss function (see Equation 2, Materials and Methods). **(D) Design concept of the chromosome-aware module.** Local selection captures within-chromosome dependencies, while global selection captures inter-chromosomal effects. Integrating both levels preserve complementary genetic dependencies. **ALT Text:** A multi-panel diagram (A-D) illustrating the CLCNet deep learning framework. Panel A shows a neural network flow including SNP selection, FC layers, and dual output tasks (Prediction and Embedding). Panel B outlines the data processing pipeline from chromosome-level splitting to nonlinear modeling and SNP filtering. Panel C depicts the contrastive learning process where embeddings and trait values from multiple samples are paired to calculate Euclidean distance and difference values for a contrastive loss function. Panel D uses horizontal bars and curved arrows to represent inter-chromosomal and intra-chromosomal interactions.

The framework integrates two key modules. First, a chromosome-aware SNP preprocessing module leverages genomic structure by grouping SNPs by chromosome and performing feature selection at both chromosome-level (local) and genome-wide (global) scales (Fig. 1 B and D). Informative SNPs were identified using LightGBM based on information gain (IG > 0). Local selection preserves within-chromosome dependencies, such as linkage disequilibrium (LD) blocks, whereas global selection captures inter-chromosomal relationships. Non-overlapping SNPs from both levels are combined to form the final input.

Second, a contrastive learning module was integrated into the multi-task framework to stabilize feature representation (Fig. 1 C). During training, samples are randomly paired at each epoch, and a composite loss is optimized, combining the phenotypic prediction loss (MSE) with a customized contrastive loss (see Equation 2 in Materials and Methods section). This loss aligns the Euclidean distance between SNP embeddings with the phenotypic differences between paired samples, encouraging phenotypically similar samples to cluster while separating dissimilar ones.

By integrating chromosome-aware feature selection with contrastive learning, CLCNet simultaneously derives accurate trait predictions and informative sample-level representations. The framework is well suited for genomic selection scenarios characterized by high-dimensional SNP data (>100,000 markers) and relatively small sample sizes (<10,000), as commonly observed in modern breeding programs.

### Superior performance of CLCNet over other models

Publicly available data from CropGS-Hub [31], a large-scale genomic selection resource that integrates genotypic and phenotypic information, were used in this study. Four representative datasets were curated, covering maize (*Zea mays*), cotton (*Gossypium hirsutum*), rapeseed (*Brassica napus*), and soybean (*Glycine max*). In total, ten agronomically important traits were analyzed, including days to tasseling (DTT), ear weight (EW), plant height (PH), flowering time (FT), oil content (OC), protein content (PC), hundred seed weight (HSW), seed length (SL), fiber strength (FStr), and fiber micronaire (FMic). These datasets exhibit substantial heterogeneity, with sample sizes ranging from fewer than 1,000 to over 8,600 individuals and raw SNP counts from ~1 million to ~30 million (Supplementary Table 1). They also span diverse genetic architectures (Supplementary Table 2; Supplementary Fig. 2), with linkage disequilibrium (LD; *r*^2^ = 0.21-0.74) and heritability (*h*^2^ = 0.49-0.99) varying widely. After LD filtering, the number of SNPs remained high, ranging from ~125,000 to over 660,000 markers, reflecting the high-dimensional, low-sample-size setting typical of genomic selection. CLCNet was benchmarked against a diverse set of baseline models, with performance evaluated on held-out test sets generated from a 3 × 10-fold cross-validation scheme.

In maize, CLCNet achieved the highest PCC across three traits (Fig. 2; Supplementary Table 3). Performance gains were statistically significant, except for DNNGP on EW (Fig. 2 B) and SVR on PH (Fig. 2 C). Among baseline methods, rrBLUP, SVR, and DNNGP were the best-performing models within the statistical, machine learning, and deep learning categories, respectively. Relative to these, CLCNet improved PCC by 0.34%, 0.46%, and 0.77% on DTT; 0.98%, 0.58%, and 0.23% on EW; 0.36%, 0.03%, and 1.15% on PH. In addition, LightGBM, DeepGS, and DNNGP showed the weakest performance on DTT, EW, and PH, respectively, where CLCNet achieved larger improvements of 1.31%, 2.76%, and 1.15%. In terms of MSE, CLCNet consistently achieved the lowest values, with statistically significant improvements (Supplementary Fig. 1 A-C; Supplementary Table 4), with only a few exceptions, consistent with the PCC results.

**Fig. 2.**
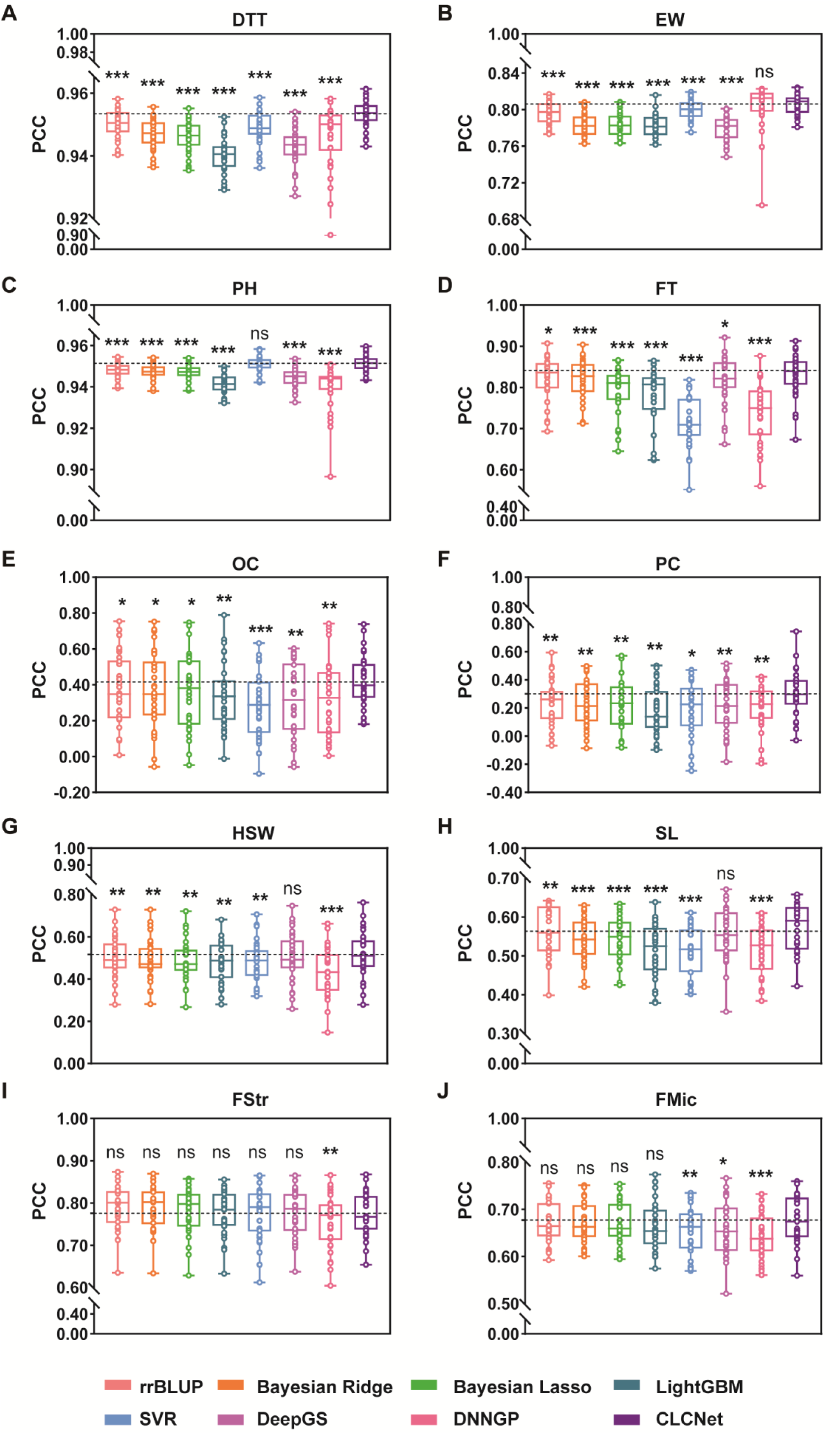
Prediction performance of CLCNet on ten traits by PCC. **(A) maize DTT, (B) maize EW, (C) maize PH, (D) rapeseed FT, (E) rapeseed OC, (F) rapeseed PC, (G) soybean HSW, (H) soybean SL, (I) cotton FStr and (J) cotton FMic.** Trait abbreviations correspond to: DTT-days to tasseling, EW-ear weight, PH-plant height, FT-flowering time, OC-oil content, PC-protein content, HSW-hundred seed weight, SL-seed length, FStr-fiber strength, and FMic-fiber micronaire. The dashed line represented the average Pearson correlation coefficient (PCC) of CLCNet. A two-tailed paired t-test with Benjamini-Hochberg false discovery rate (FDR) correction was used to compare CLCNet with each model. *** denoted *p* < 0.001, ** denoted *p* < 0.01, * denoted *p* < 0.05, and ns indicated no significant difference. **ALT Text**: A grid of ten box plots (A-J) comparing the prediction performance (PCC) of CLCNet against other models across various crops and traits. Each panel contains multiple colorful box plots with overlaid individual data points, representing different models. A horizontal dashed line indicates the mean performance of CLCNet for reference. Statistical significance markers (asterisks or “ns”) are placed above each model’s box plot to indicate performance differences relative to CLCNet. A color-coded legend at the bottom identifies the specific models compared.

A similar pattern was observed in rapeseed. CLCNet ranked first in PCC for three traits (Fig. 2 D-F; Supplementary Table 3). Compared with rrBLUP, the second-best model, CLCNet improved PCC by 0.62% for FT, 4.35% for OC, and 6.53% for PC. Compared with other models, including DeepGS, DNNGP, LightGBM, and SVR, the gains were even greater: for FT, PCC increased by 1.16%, 9.09%, 4.65%, and 11.44%; for OC, gains reached 10.02%, 8.98%, 7.59%, and 13.31%; and for PC, improvements were 8.37%, 10.13%, 12.19%, and 10.12%. MSE results were consistent with the PCC evaluations (Supplementary Fig. 1 D-F; Supplementary Table 4), although rrBLUP, Bayesian Ridge, and Bayesian Lasso exhibited comparable errors to CLCNet.

In soybean, CLCNet demonstrated robust advantages for HSW and SL, achieving significantly higher PCC values, together with lower MSE values (Fig. 2 G-H; Supplementary Fig. 1 G-H; Supplementary Table 3). Specifically, for HSW, CLCNet exceeded the second-best model, DeepGS, by 1.35% and outperformed the lowest performing DNNGP by 8.41%. For SL, PCC improvements relative to the second-best rrBLUP and lowest-performing LightGBM were 0.80% and 5.90%, respectively.

In cotton, performance differences among models were generally smaller (Fig. 2 I; Supplementary Fig. 1 I; Supplementary Table 3). For FMic, CLCNet achieved PCC values of 67.70%, comparable to rrBLUP, Bayesian Ridge, Bayesian Lasso, and LightGBM, with no statistically significant differences. However, CLCNet significantly outperformed SVR, DeepGS, and DNNGP, with PCC improvements of 2.33%, 1.86%, and 3.56%, respectively. Consistently, MSE analysis showed that CLCNet had significantly lower prediction errors than SVR and DNNGP (Supplementary Table 4).

Increasing cross-validation repetitions from 3 to 10 did not alter model rankings, confirming the robustness of the results (Supplementary Table 3; Supplementary Table 4). Variance decomposition further indicated that performance variability was primarily driven by data partitioning rather than random initialization.

Overall, CLCNet achieved consistent improvements in PCC (0.34%–12.19%) and reduced MSE across traits and species (Supplementary Table 3; Supplementary Table 4). Performance gains were more pronounced for traits with moderate LD (*r*^2^ = 0.21–0.36) and high heritability (*h*^2^ > 0.66), such as those in maize, rapeseed, and soybean (Supplementary Table 2; Supplementary Fig. 2). In contrast, for cotton traits characterized by higher LD (*r*^2^ = 0.74) and lower heritability (*h*^2^ < 0.50), model differences were smaller, suggesting that largely additive genetic architectures are already well captured by linear approaches. Notably, CLCNet maintained robust performance under these conditions.

### Strong feature extraction capability of the Chromosome-aware module

Feature extraction is critical in GP for identifying informative SNPs, reducing redundancy, and improving model accuracy [32]. We compared chromosome-aware (CA) module with two widely used feature selection methods: genome-wide association study (GWAS) and principal component analysis (PCA). For GWAS, SNPs with *p* < 0.01 were selected, whereas for PCA, principal components explaining more than 98% of the variance were retained as input. All eight models were trained using each strategy and evaluated by PCC and MSE.

Across traits, CLCNet with the CA module consistently achieved higher PCC and lower MSE than PCA, with statistically significant improvements. Specifically, PCC gains of CA over PCA ranged from 1.40% (PH) to 18.76% (PC), and over GWAS from 0.20% (PH) to 4.06% (HSW) (Fig. 3 B; Supplementary Table 5). MSE results showed similar trends, with CA yielding lower errors across most traits (Supplementary Table 6).

**Fig. 3.**
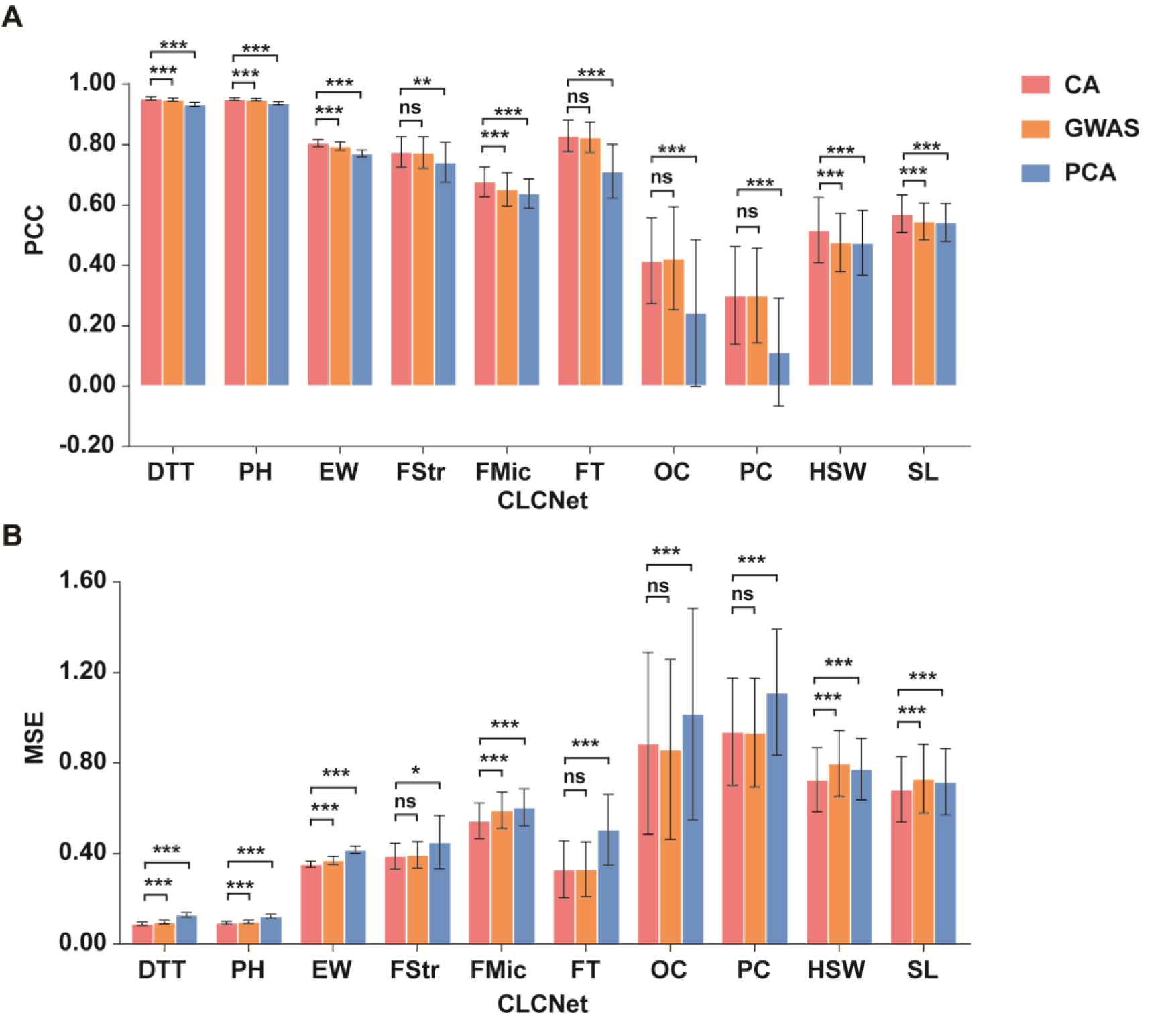
Performance comparison of feature selection methods based on PCC and MSE in CLCNet. **(A) The predictive performance of CLCNet using chromosome-aware (CA), GWAS-based, and PCA-based feature selection methods was evaluated.** *** denoted *p* < 0.001, ** denoted *p* < 0.01, * denoted *p* < 0.05, and ns indicated no significant difference. **ALT Text:** Two clustered bar charts (A and B) comparing three feature selection methods: CA (red), GWAS (orange), and PCA (blue) across ten traits (DTT, PH, EW, etc.). Each trait group contains three colored bars with error bars and statistical significance brackets (asterisks or “ns”). A legend on the right identifies the colors for CA, GWAS, and PCA.

Similar patterns were observed across other models (rrBLUP, Bayesian Ridge, Bayesian Lasso, and DeepGS). PCC gains of CA (e.g., over PCA, GWAS) ranged from approximately 0.11% (Bayesian Lasso for PH using GWAS) to 34.54% (DeepGS for FStr using PCA) (Supplementary Fig. 3-6; Supplementary Table 5). Compared with no feature selection (green bars), CA significantly improved PCC in 7–10 traits, with increases ranging from 0.07% (Bayesian Ridge for PH) to 5.50% (Bayesian Ridge for PC).

Model-dependent differences were evident. In LightGBM, CA showed no significant advantage over the no-selection setting or GWAS, likely because CA is derived from LightGBM feature importance (Supplementary Fig. 7). However, CA still significantly outperformed PCA in eight traits (PH, PC, FStr, FMic, FT, OC, HSW, and SL), with gains ranging from 0.35% to 10.27%, highlighting the limitations of linear projection-based dimensionality reduction in tree-based models (Supplementary Table 5).

In SVR, GWAS showed strong compatibility, and CA outperformed GWAS significantly only for FT (Supplementary Fig. 8; Supplementary Table 5-6). Nevertheless, CA improved PCC over no-selection in nine traits, with gains ranging from 0.13% (PH) to 8.60% (OC), and outperformed PCA in five traits (DTT, PH, EW, FStr, and FMic), with gains up to 20.14% (PH).

In DNNGP, CA showed significant advantages over PCA in only three traits (FStr, FMic, and FT), with PCC increases of 1.56%, 3.61%, and 3.54%, respectively (Supplementary Fig. 9; Supplementary Table 5-6). These results reflect the strong coupling between DNNGP and PCA-derived features.

We further assessed overall performance by ranking models based on mean PCC (Fig. 4). CLCNet ranked first in two traits (DTT and EW) and no lower than sixth (FMic) in any trait. Notably, the top four positions were exclusively occupied by CLCNet or CA-based models, underscoring the effectiveness of the CA design in achieving reliable genomic prediction.

**Fig. 4.**
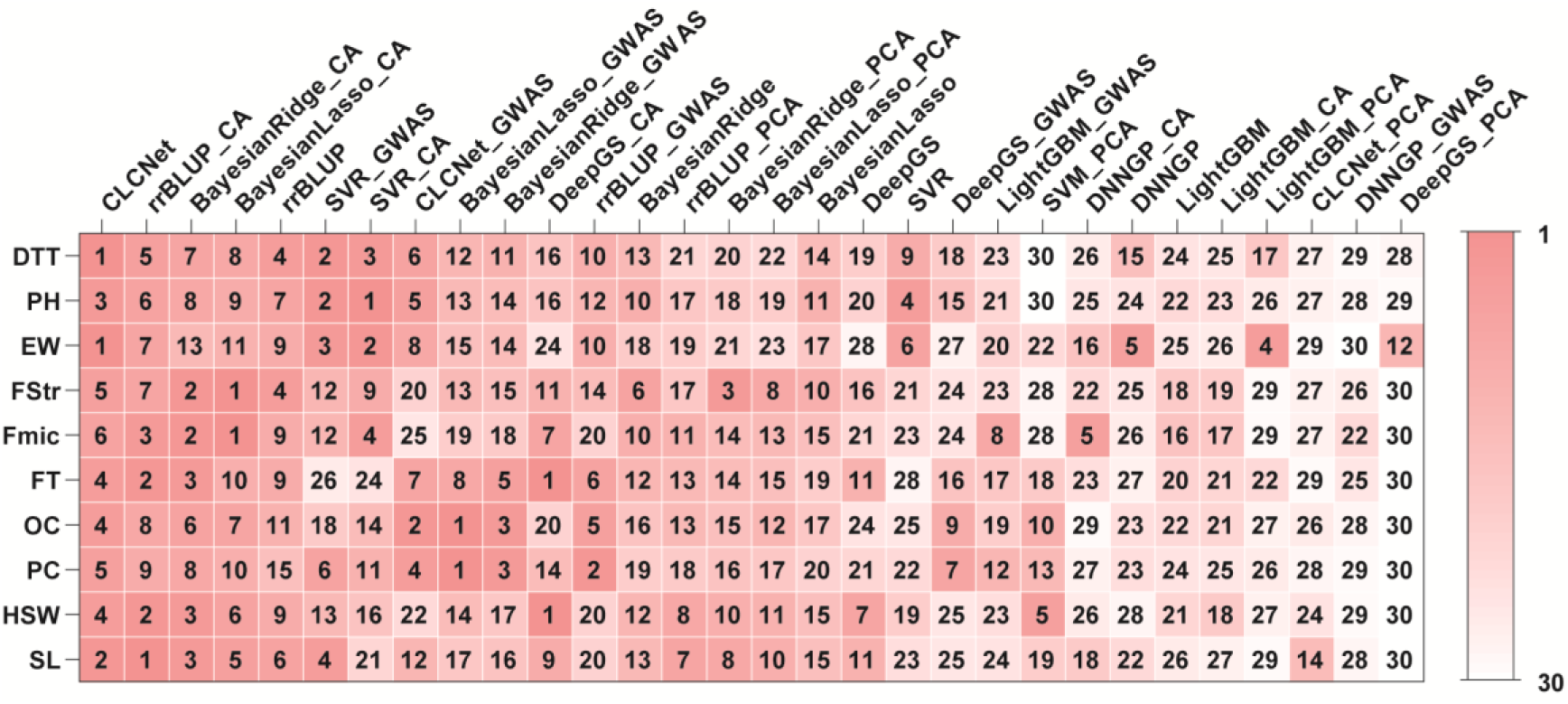
Comparative ranking of feature selection strategies across models and traits. Models were evaluated under four different selection strategies: no-selection, CA selection, GWAS selection, and PCA selection. For each trait, models were ranked from 1 to 30 based on mean PCC values averaged over three independent repetitions of randomly stratified ten-fold cross-validation. **ALT Text:** A heatmap displaying the ranking of 30 different combinations of models and feature selection strategies across 10 traits (shown as rows). The x-axis labels at the top list the model-strategy combinations (e.g., CLCNet, rrBLUP_CA, BayesianLasso_GWAS). Each cell contains a rank number from 1 to 30, color-coded with a red-to-white gradient scale. Darker red cells indicate higher rankings (approaching 1), while lighter or white cells indicate lower rankings (approaching 30). A color scale bar on the right side of the heatmap illustrates the gradient from rank 1 to 30.

Overall, the CA module demonstrated consistent advantages across models and traits. PCC gains over PCA reached up to 34.54%, while improvements over GWAS were smaller but stable (0.11%-8.39%). Ranking analysis further confirmed this advantage (Fig. 4, first four columns).

### Effectiveness of independently designed modules in CLCNet

To evaluate the contributions of the chromosome-aware feature selection and contrastive learning modules, we conducted ablation studies using four CLCNet variants (Supplementary Fig. 10). Specifically, CLCNet_1 removed the genome-wide SNP feature selection (global) while retaining chromosome-level SNP selection (local). CLCNet_2 further excluded the contrastive learning module from CLCNet_1, reducing the model to a single-task setting. In contrast, CLCNet_3 retained only the global selection by removing local selection. Finally, CLCNet_4 further removed the contrastive learning from CLCNet_3, yielding a minimal single-task baseline.

Across traits, no significant differences were observed between CLCNet and CLCNet_1, indicating that removal of the global selection alone had a limited impact on predictive performance when local selection was retained (Fig. 5; Supplementary Table 7-8). In contrast, removing local selection led to substantial performance degradation. Significant differences between CLCNet and CLCNet_3 were observed (Fig. 5; Supplementary Fig. 11; Supplementary Table 7-8), with PCC reductions ranging from 0.23% (PH) to 14.22% (PC). Similar patterns were observed in CLCNet_4, with gains ranging from 0.36% (PH) to 14.22% (OC), confirming the critical role of chromosome-aware local selection.

**Fig. 5.**
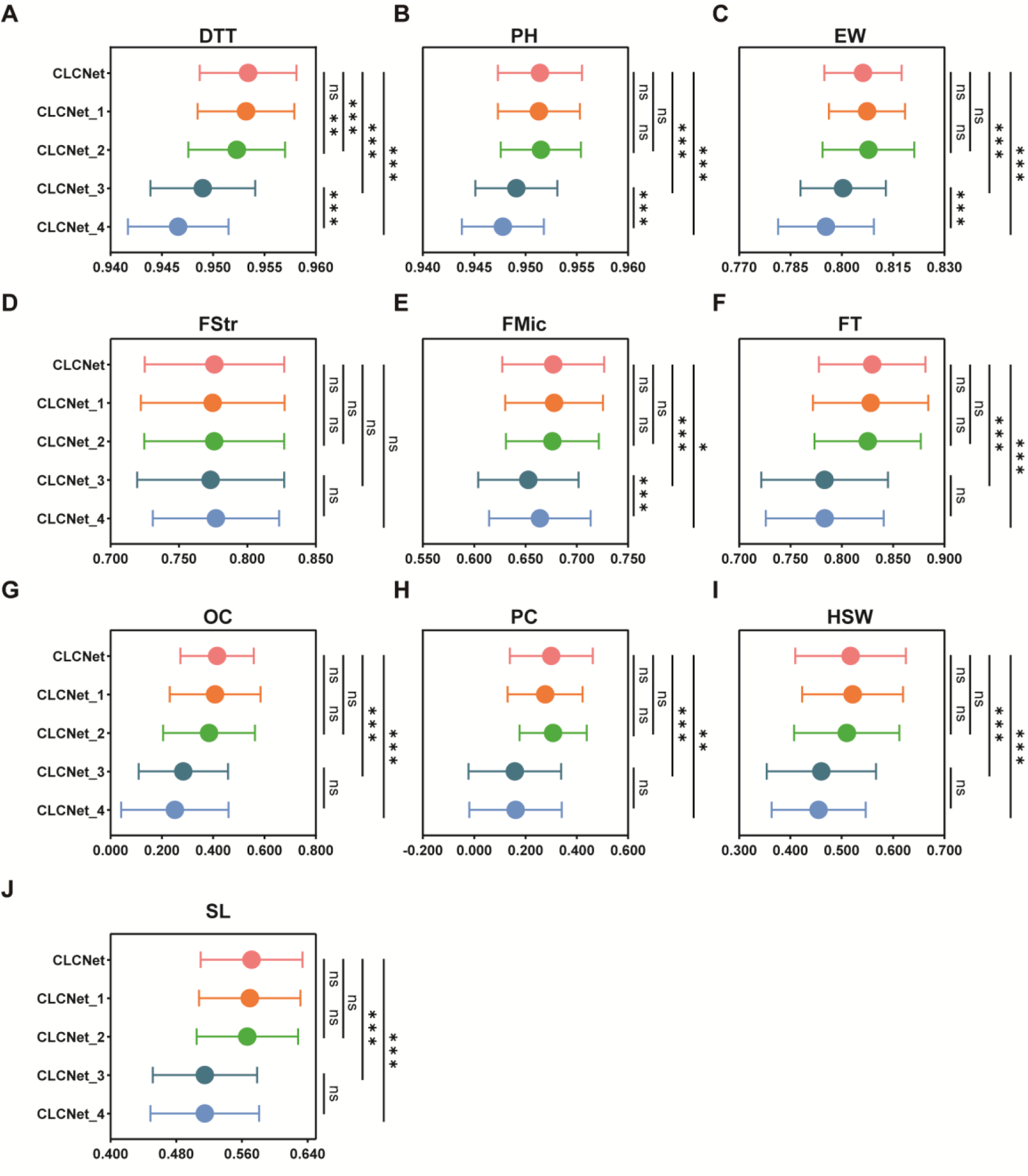
Performance of CLCNet’s core modules on PCC. **(A) maize DTT, (B) maize PH, (C) maize EW, (D) cotton FStr, (E) cotton FMic, (F) rapeseed FT, (G) rapeseed OC, (H) rapeseed PC, (I) soybean HSW and (J) soybean SL.** Statistical significance was assessed using a two-tailed paired t-test with Benjamini–Hochberg false discovery rate (FDR) correction. *** denoted *p* < 0.001, ** denoted *p* < 0.01, * denoted *p* < 0.05, and ns indicated no significant difference. **ALT Text:** A grid of ten dot plots (A-J) showing the Pearson correlation coefficient (PCC) for the CLCNet model and four variants (CLCNet_1 to CLCNet_4). In each plot, the y-axis lists the model versions, and the x-axis shows the PCC value range. Each data point is represented by a colored circle (red, orange, green, teal, and blue) with horizontal error bars indicating the standard deviation. On the right side of each plot, vertical bars with statistical symbols (asterisks or “ns”) indicate the results of significance tests between the full model and its variants.

The contribution of contrastive learning was more moderate and trait-dependent. Comparisons of CLCNet_1 vs. CLCNet_2 and CLCNet_3 vs. CLCNet_4 showed significant performance gains for DTT, PH, and EW (Fig. 5 A, F, I; Supplementary Tables 7-8), underscoring the module’s capacity to refine phenotypic differentiation. For other traits, including FT, OC, HSW, SL, and FMic, its removal consistently reduced performance, albeit without statistical significance. These results suggest that contrastive learning provides a consistent but incremental benefit.

Overall, the ablation analysis reveals a clear hierarchy of contributions: local selection has the largest impact, followed by contrastive learning, while global selection contributes relatively little.

### Chromosomal structure-guided SNP selection in the Chromosome-aware module

Our ablation analyses demonstrated that chromosome-level feature selection (local) contributed most to model performance. We hypothesized that this superior performance stems from the explicit integration of chromosomal structural priors. To validate this, we performed a perturbation experiment that disrupted chromosomal structure while preserving SNP composition (Supplementary Fig. 10 F). Specifically, SNP positions within each chromosome were randomly permuted, breaking the original positional order while maintaining the same number of SNPs per chromosome. This manipulation removed biologically meaningful positional context without altering allele frequencies or overall feature dimensionality. By comparing the predictive performance of CLCNet before and after chromosomal shuffling, we directly assessed whether the CA module relies on chromosomal structural information rather than merely on the presence of informative SNPs.

Consistent with our hypothesis, chromosomal shuffling led to a systematic decline in predictive accuracy. Nine traits showed significant PCC reductions, ranging from 0.12% (PH) to 9.88% (OC) (Fig. 6; Supplementary Table 9). MSE exhibited consistent increases across the same traits (Supplementary Fig. 12; Supplementary Table 10), confirming that performance degradation was driven by disruption of structural information rather than random variation.

**Fig. 6.**
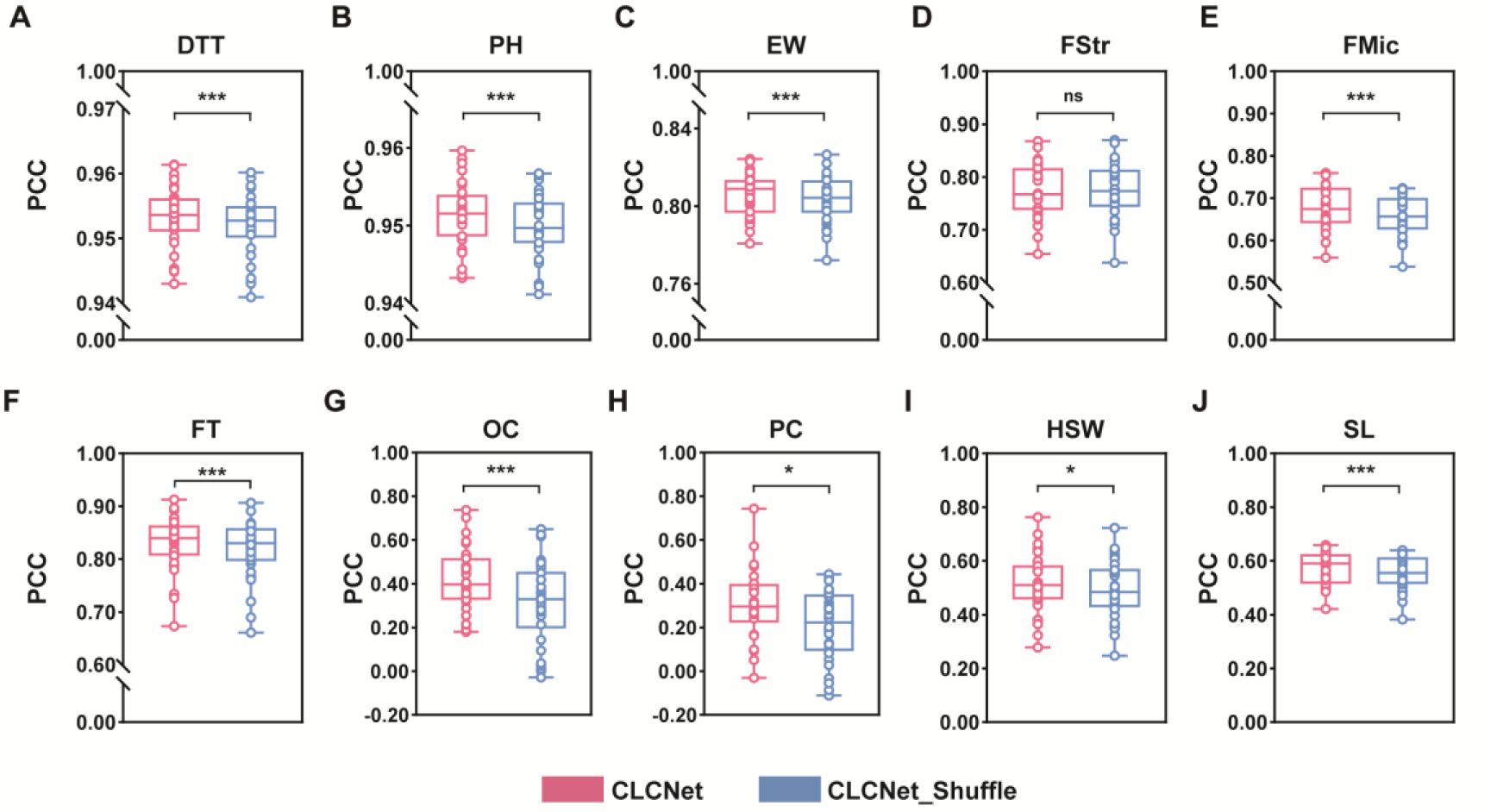
Model PCC performance under SNP position perturbation. **(A) maize DTT, (B) maize PH, (C) maize EW, (D) cotton FStr, (E) cotton FMic, (F) rapeseed FT, (G) rapeseed OC, (H) rapeseed PC, (I) soybean HSW and (J) soybean SL.** CLCNet was compared with its shuffled variant (CLCNet_Shuffle) using a two-tailed paired t-test with Benjamini–Hochberg false discovery rate (FDR) correction. *** denoted *p* < 0.001, ** denoted *p* < 0.01, * denoted *p* < 0.05, and ns indicated no significant difference. **ALT Text:** A grid of ten paired box plots (A-J) comparing the prediction performance (PCC) of the original CLCNet model (pink) and its shuffled variant, CLCNet_Shuffle (blue), across different traits. Each plot features y-axis breaks to highlight the data distribution and the relevant PCC range. The box plots include overlaid individual data points, showing the median and quartiles for both models. Horizontal brackets with statistical significance markers (asterisks or “ns”) are positioned above each pair of boxes to indicate the impact of SNP position perturbation on model performance. A legend at the bottom defines the colors for CLCNet and CLCNet_Shuffle.

Collectively, these results highlighted the critical role of SNP positional information in chromosome-aware modeling. The pronounced decline provides strong evidence that CLCNet effectively captures structured genomic signals.

### Evaluating SHAP-Based SNP importance against GWAS results

GWAS has been widely used to identify genetic variants associated with complex traits [33]. However, by testing SNPs individually, it overlooks SNP–SNP interactions and nonlinear genotype–phenotype relationships. Deep learning models such as CLCNet can complement these limitations. We applied SHAP (SHapley Additive exPlanations) [34] to estimate SNP importance, and compared these scores with GWAS p-values and effect sizes to facilitate gene discovery and assess the biological relevance of the identified SNPs.

GWAS were performed for three maize traits using 125,820 SNPs (Supplementary Table 1), while the CA module selected approximately 21,315 SNPs. Many SNPs prioritized by CLCNet corresponded to relatively lower −logDD(p) values in GWAS Manhattan plots (Fig. 7 A; Supplementary Fig. 13 A and 14 A). SHAP profiles of selected SNPs showed peak patterns broadly consistent with GWAS signals (Fig. 7 C; Supplementary Fig. 13 C and 14 C). These results suggested that CLCNet captures overlapping yet distinct genetic information.

**Fig. 7.**
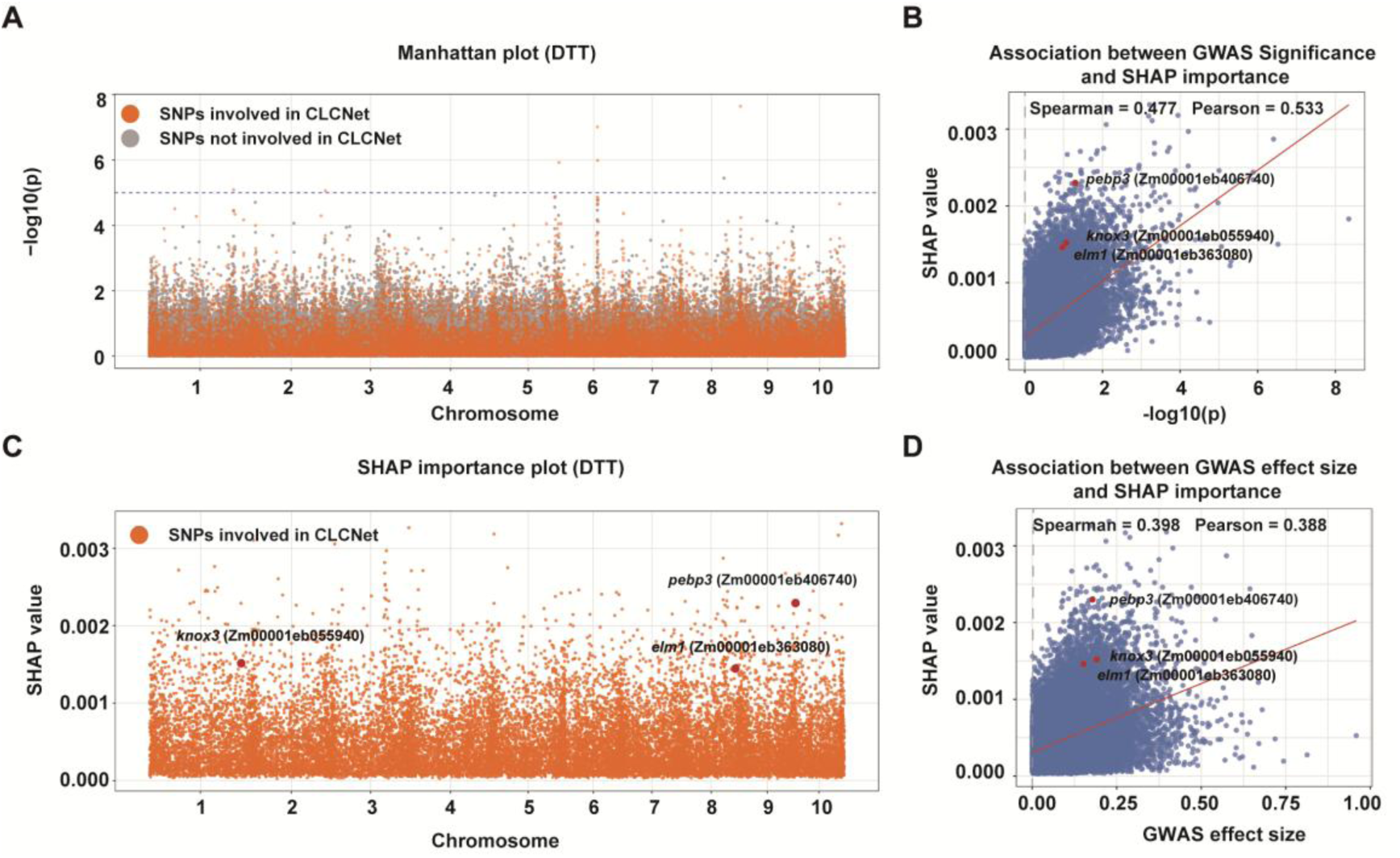
Relationship between SHAP-based SNP importance derived from CLCNet and GWAS for DTT. (A) Manhattan plot of GWAS results for DTT. (B) Relationship between GWAS association significance (–log□□(p)) and SNP importance scores derived from CLCNet using SHAP. (C) Distribution of SHAP-based SNP importance scores generated by CLCNet. (D) Relationship between GWAS-estimated effect size and SNP SHAP importance scores by CLCNet. **ALT Text:** A composite figure with four panels (A-D) illustrating the relationship between SHAP-derived SNP importance and GWAS results for the DTT trait. Panels A and C are Manhattan-style plots showing the distribution of SNPs across 10 chromosomes; Panel A colors SNPs based on their involvement in CLCNet, while Panel C focuses on SHAP importance distribution. Panels B and D are scatter plots showing positive correlations between SHAP values and GWAS statistics, featuring red regression lines and specific gene labels such as *pebp3*, *knox3*, and *elm1* highlighted in red. Correlation values (Spearman and Pearson) are displayed at the top of the scatter plots.

Correlation analysis further supported this relationship. Spearman and Pearson correlations between SHAP importance and GWAS statistics across the three traits ranged from 0.3981 to 0.5092 and from 0.3885 to 0.5733 (Fig. 7 B and 7 D; Supplementary Fig. 13 B and 13 D; Supplementary Fig. 14 B and 14 D), respectively. Based on the top 5% SHAP-ranked SNPs, several biologically relevant candidate genes were identified (Supplementary Table 11). For DTT, the model highlighted *pebp3* (Zm00001eb406740), a member of the PEBP family known to regulate flowering time [35]. It also identified *knox3* (Zm00001eb055940), which is involved in meristem-related development [36], and *elm1* (Zm00001eb363080), associated with cell elongation [37]. For PH, key candidates included *ys1* (Zm00001eb249020), which controls iron uptake and affects plant growth [38], and *rs2* (Zm00001eb029300), a MYB transcription factor regulating leaf development and shoot architecture [39]. In addition, *pebp6* (Zm00001eb197000) may suggest a potential link between flowering-related and plant height, given the known role of PEBP family genes [35]. For EW, the identification of *bd1* (Zm00001eb330200), a critical regulator of spikelet meristem identity [40]. The gene *tan1* (Zm00001eb286860), involved in cell division orientation, may indirectly influence kernel development and ear architecture [41]. These results suggest that CLCNet recapitulates biologically relevant GWAS signals while capturing additional information beyond conventional approaches.

## Discussion

CLCNet adopts a multi-task learning (MTL) framework, which fundamentally distinguishes it from single-task learning, multi-modal learning, and transfer learning approaches. Prior work has shown that MTL enhances performance on related tasks by leveraging shared representations [42]. In this study, we further integrate contrastive learning into the MTL framework to strengthen representation learning by explicitly modeling inter-sample variation in genotype–phenotype relationships. Our results indicate that contrastive learning acts as a synergistic component, as evidenced by performance improvements in maize (e.g., ablation comparisons between CLCNet_1 vs. CLCNet_2 and CLCNet_3 vs. CLCNet_4). In GP, high-dimensional features coupled with limited sample sizes often lead to model instability and heightened sensitivity to stochastic noise, particularly when encountering samples with divergent phenotypes or genotypes. Contrastive learning mitigates these effects by explicitly modeling individual-level relationships-effectively clustering congruent samples while dissociating divergent ones. Notably, the effectiveness of this appears dependent on sample size. Gains were most pronounced in the largest dataset (n = 8,652; Supplementary Table 1), which far exceeds the smaller cohorts (n = 991-2,795). This larger sample pool provides a richer and more diverse set of pairwise relationships under the contrastive loss.

The chromosome-aware module proposed in this work offers a biologically informed feature selection strategy. In high-dimensional genomic datasets where SNP features vastly outnumber phenotypic samples, dimensionality reduction is essential to address the “curse of dimensionality” [32]. Feature selection reduces noise and redundancy while preserving informative genetic signals [43]. However, conventional GWAS-based feature selection methods primarily rely on linear models to evaluate associations between individual SNPs and phenotypes. Similarly, commonly feature selection methods such as principal component analysis (PCA) [20], maximal information coefficient (MIC) [21], and Pearson-collinearity selection (PCS) [44] typically do not explicitly account for grouped interactions. CLCNet implements an explicit chromosome-based grouping strategy that directly leverages biological chromosome structure. Our results demonstrate that local, within-chromosome SNP selection plays a decisive role in improving prediction accuracy. The substantial performance decline following random permutation of SNP positions provides strong evidence that the chromosome-aware module captures biologically meaningful positional and structural information beyond individual SNP effects.

The biological relevance of CLCNet is further corroborated by comparisons between SHAP-derived SNP importance scores and GWAS results. Moderate yet consistent correlations indicate that CLCNet recovers established GWAS signals while identifying additional complementary genetic information. The incomplete concordance between methods reflects their fundamental differences: GWAS evaluates independent marginal linear effects, whereas CLCNet captures nonlinear interactions among SNPs. Consequently, SNPs with modest marginal effects in GWAS may attain high importance in CLCNet due to their involvement in complex interaction patterns. Consistent with this, among the top 5% SHAP-ranked SNPs, several known candidate genes were identified (Supplementary Table 11), including *pebp3*, *knox3*, *elm1* (DTT), *ys1*, *rs2*, *pebp6* (PH), and *bd1*, *tan1* (EW), which are typically not prioritized by GWAS (Fig. 7 B and D; Supplementary Fig. 13 B and D; Supplementary Fig. 14 B and D).

Although the PCC gains over strong baseline models are modest (approximately 1% for some traits), CLCNet consistently improves Spearman’s rank correlation across traits (Supplementary Table 12). Given that breeding decisions primarily depend on ranking individuals, these gains are practically meaningful. In maize, improvements range from 0.32% to 3.20%, whereas in rapeseed and soybean they are considerably larger, reaching up to 13.70% and 7.24%, respectively. In a population of 10,000 individuals, even a 1% increase can shift dozens to over a hundred individuals into the top selection set. Consistent with the breeder’s equation [45], genetic gain is proportional to selection accuracy, and such gains accumulate over cycles in long-term breeding programs.

However, this study still has several limitations. First, the proposed dual-pathway architecture of CLCNet required relatively long training time and inference time (Supplementary Table 13). For maize data comprising 8,652 samples and 125,820 SNPs, CLCNet required 134–795 s per trait, compared with 74-373 s for DNNGP, 174-216 s for DeepGS, 0.08-106 s for machine learning models (SVR, LightGBM), and 108-271 s for statistical models (rrBLUP, Bayesian Ridge, Bayesian Lasso). Second, CLCNet did not include comparisons with a wider range of methods, particularly recently developed deep learning models such as VMGP [46] and MeNet [47]. Third, the generalization of CLCNet across diverse population structures and environmental conditions was not explicitly assessed. In practical breeding, population heterogeneity and genotype-by-environment interactions inevitably influence model performance. While cross-validation across species and traits suggests robustness, this does not fully replicate real-world deployment scenarios. Future work will address these limitations.

In summary, we introduce CLCNet, a novel deep learning model for plant genomic prediction that integrates contrastive learning and chromosome-aware modules for the first time [48]. CLCNet consistently outperforms linear models, machine learning models, and classical deep learning models. Collectively, our results demonstrate that CLCNet provides a robust and scalable solution for high-dimensional, low-sample-size GP, with the potential to accelerate genetic gain in plant breeding.

## Materials and methods

### Data description and preprocessing

Four representative datasets were obtained from the CropGS-Hub database [31]. Genotypes were encoded as 0, 1, or 2, with missing single-nucleotide polymorphisms (SNPs) imputed using a constant value of 3. This strategy enables the model to distinguish missing entries from observed genotypes without invoking assumptions regarding allele frequencies. Samples lacking phenotypic records were removed; no further filtering was applied. Linkage disequilibrium (LD) pruning was performed using PLINK (v1.9) [49] with parameters --indep-pairwise 100 10 0.2 to reduce data dimensionality. The final datasets comprised maize (*Zea mays*) from GSTP001, cotton (*Gossypium hirsutum*) from GSTP010, rapeseed (*Brassica napus*) from GSTP013, and soybean (*Glycine max*) from GSTP014, encompassing a total of 10 traits. A summary of the preprocessed datasets was provided in Supplementary Table 1.

### The architecture of CLCNet

CLCNet is a multi-task deep learning model implemented in PyTorch (v2.1.0; https://pytorch.org/) that accepts genomic SNP data as input and outputs phenotypic predictions as the primary task alongside low-dimensional feature embeddings as an auxiliary task. The backbone adopted a dual-pathway architecture. One pathway consisted of three nonlinear fully connected layers with ReLU activation functions, with layer dimensions of {4096,2048, 1024}. The other pathway comprised a single linear fully connected layer with an output dimension of 1024. The outputs of the two pathways were summed at the 1024-dimensional level and subsequently fed into both the primary and auxiliary task branches. Each branch was composed of two nonlinear fully connected layers. For the primary task, the layer dimensions were {4096, 2048,1}, producing the final phenotypic prediction, whereas for the auxiliary task, the dimensions were {4096,2048,1024}, generating low-dimensional feature embeddings.

### Chromosome-aware module

A chromosome-aware module was incorporated as a preprocessing step for genomic data. SNPs were first grouped into chromosomal level according to their genomic positions. Local feature selection was then performed within each chromosome to identify informative SNPs (referred to as local SNPs). In parallel, global feature selection was applied across the entire genome to capture genome-wide informative SNPs (referred to as global SNPs). The selected local and global SNPs were concatenated, with duplicate SNPs removed, to form the final input. Feature selection was conducted using the LightGBM algorithm, a tree-based method capable of modeling nonlinear relationships among SNPs [50]. SNPs with positive information gain (IG > 0) were retained, while those with zero information gain (IG = 0) were discarded.

### Contrastive learning module

A contrastive learning module was introduced through a customized training strategy. During each training epoch, a random pairing mechanism was used when loading samples, such that each sample was paired with another randomly selected sample. The paired samples were fed into the model simultaneously, and both their phenotypic values and low-dimensional feature embeddings were predicted. Model optimization was guided by three loss components: the mean squared error (MSE) between the predicted and true phenotypic values for each of the two input samples (Equation 1), and a contrastive loss based on the Euclidean distance between the embeddings of the paired samples and the difference in their phenotypic values (Equation 2).

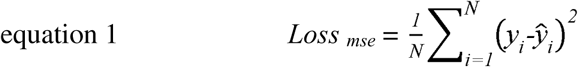

Here, *N* is the number of samples. y*_i_* is the true value for the *i*-th observation. *ŷ_i_* is the predicted value for the *i*-th observation.

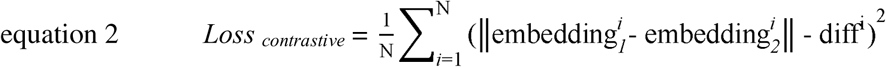

Here, *N* is the number of samples. ||embedding*^i^*_1_ - embedding*^i^*_2_|| denotes the Euclidean distance between the *i*-th pair of embeddings. diff*^i^* is the phenotypic difference for the *i*-th pair of samples.

### Model comparison and cross-validation

The predictive performance of CLCNet was evaluated against models on benchmark datasets, including three classical statistical methods, rrBLUP [7], Bayesian Ridge and Bayesian Lasso [51]; two machine learning algorithms, LightGBM [52] and SVR [53]; and two deep learning frameworks, DeepGS [17] and DNNGP [20]. Classical methods were implemented using the R packages rrBLUP (v4.6.2) and BGLR (v1.1.3). For machine learning, LightGBM was implemented with Python’s lightgbm package (v3.3.5), while SVR utilized Python’s scikit-learn library (v0.24.2). DeepGS utilized a convolutional neural network (CNN) architecture comprising input, convolutional, max-pooling, dropout, fully connected, and output layers, with dropout rates of 0.2, 0.1, and 0.05 applied at various stages to mitigate overfitting. DNNGP adopted a featuring three convolutional layers, batch normalization, dropout, a flattening layer, a dense layer, and an output layer. Principal components accounting for over 98% of the variance were extracted from the genomic data using PLINK (v1.9) as input features.

A 3 × 10-fold cross-validation (CV) strategy was adopted. Specifically, ten-fold CV was repeated three times using different random data partitions (random seed 42, 84 and 162), resulting in a total of 30 independent training and testing processes. In each fold, the training data were further randomly split into an internal training set (80%) and an internal validation set (20%), while the test set was strictly held out for final evaluation. All models were trained exclusively on the internal training set. Feature selection strategies, including chromosome-aware selection (CA), Genome-wide association study (GWAS) and principal component analysis (PCA), were conducted only on the internal training set, and subsequently applied to the internal validation and test sets. The test sets were used exclusively for final performance assessment and were not involved in any stage of model training, feature selection, or hyperparameter optimization. Phenotype values were standardized using z-score normalization based solely on the mean and variance computed from the internal training set to prevent data leakage. The same normalization parameters from the internal training set were then applied to the internal validation and test sets.

### Hyperparameter optimization

Hyperparameter tuning was performed using grid search for both machine learning and deep learning models. For LightGBM, the number of leaves {num_leaves: 31, 63}, the maximum tree depth (max_depth: 5, 10, −1), the learning rate {learning_rate: 0.01, 0.05, 0.1}, and the number of boosting rounds (n_estimators: 100, 200, 500) were systematically explored. For SVR, the regularization parameter {C: 0.1, 1, 10, 100}, the epsilon-insensitive loss parameter {epsilon: 0.01, 0.1, 0.5}, the kernel coefficient {gamma: ‘scale’, ‘auto’}, were evaluated with the RBF kernel. For deep learning models, including CLCNet, DeepGS, and DNNGP, the key hyperparameters considered included the learning rate, batch size, and number of training epochs, with candidate values of {0.01, 0.001, 0.0001}, {16, 32, 64}, and {20, 50, 100}, respectively.

The optimal hyperparameter combination for each model was automatically selected based on the highest Pearson correlation coefficient (PCC) on the internal validation set. To reduce the computational cost of hyperparameter tuning, the grid search was performed using only the internal training and validation sets of the first fold from the first (random seed 42) of three independent 10-fold cross-validation runs. Detailed hyperparameter optimization results were provided in Supplementary Table 14-18.

### Feature selection comparison

To benchmark the effectiveness of chromosome-aware (CA) module, we compared it with two widely used feature selection strategies: genome-wide association study (GWAS) and principal component analysis (PCA). For GWAS filtering, SNPs were scored using a mixed linear model implemented via GCTA [54], and markers with significant associations (*p* < 0.01) were retained. For PCA, dimensionality reduction was performed using the *sklearn.decomposition.PCA* function in Python. Consistent with DNNGP, principal components explaining more than 98% of the total variance in the genotype data were retained as model input. Each of models was evaluated by three strategies.

### SNP position perturbation

The input genotype data were represented as an *n*×*m* matrix, where *n* denotes the number of samples and *m* denotes the number of single-nucleotide polymorphisms (SNPs), ordered according to their physical positions along the genome. A single global random permutation was applied to the *m* SNP columns, while keeping the permutation identical across all samples. This operation preserved the original genotype values for each sample but completely disrupted the positional ordering of SNPs. After perturbation, the input matrix retained the same *n*×*m* dimensionality as the original data.

### SHAP-Based SNP importance analyzes

CLCNet predictions were interpreted using SHAP (SHapley Additive exPlanations) values [34]. For each trait in maize dataset, the mean absolute SHAP value of each SNP was computed. GWAS were conducted to evaluate the association between individual SNPs and the phenotype, and Manhattan plots were generated to visualize genome-wide association signals. SHAP importance scores were compared with GWAS *p*-values (–log10 transformed) and estimated effect sizes.

The raw VCF file of maize genotypes was first converted from AGPv3 coordinates to the B73 RefGen_v4 assembly using CrossMap (version 0.7.0) [55], and subsequently mapped to the Zm-B73-REFERENCE-NAM-5.0 genome coordinates. The top 5% of SNPs ranked by SHAP importance were selected for gene annotation. SNPs were mapped to genes using a ± 100 kb window, consistent with maize Linkage disequilibrium (LD) decay.

### Heritability estimation

Narrow-sense heritability (□^2^) of each trait was estimated using the genomic restricted maximum likelihood (GREML) method implemented in GCTA [54]. A genomic relationship matrix (GRM) was constructed based on genome-wide SNP. The variance components were estimated using a linear mixed model, and heritability was calculated as the proportion of genetic variance over the total phenotypic variance.

### Linkage disequilibrium decay analysis

Linkage disequilibrium (LD) decay was analyzed using PLINK. Pairwise LD between SNPs was measured using the squared correlation coefficient (*r*^2^). Specifically, LD was calculated for SNP pairs within a maximum distance of 100 kb. SNP pairs were grouped into distance bins based on their physical distance, and the average *r*^2^ value was computed for each.

### Statistical Analysis

Predictive performance was evaluated using 30 independent test sets obtained from 3 times 10-fold cross-validation and measured by the Pearson correlation coefficient (PCC) and mean squared error (MSE). Two-tailed paired t-tests were conducted to compare CLCNet with each model, including the baseline and the different CLCNet versions, with pairings defined over identical cross-validation splits. All *p*-values were adjusted using the Benjamini-Hochberg false discovery rate (FDR) correction. A same approach was used to evaluate feature selection methods, comparing GWAS and PCA with CA. All experiments were conducted on a server equipped with two Intel Xeon® Gold 6330 CPUs, 512 GB of system memory, and an NVIDIA GeForce RTX 3090 GPU.

## Supplemental information

Supplementary information is available at Briefings in Bioinformatics online.

## Acknowledgements

We would like to express our gratitude to Prof. Fei Lu, Prof. Ni Jiang, Postdoc Dr. Zeyu Zhang and Postdoc Dr. Jiachao Xu, from the Institute of Genetics and Developmental Biology, Chinese Academy of Sciences, for their professional and insightful discussions. We also thank Dr. Shihao Shao from Peking University Health Science Center for his assistance with the deep learning model. In addition, we are grateful to Prof. Jianxiao Liu from Huazhong Agricultural University for their guidance. We are especially grateful to Prof. Shichao Jin from Nanjing Agricultural University for his thoughtful suggestions and helpful discussions regarding the manuscript.

## Data availability

All datasets we used is accessible at https://iagr.genomics.cn/CropGS/#/Datasets. The scripts are available on GitHub (https://github.com/SuppurNewer/CLCNet).

