## Supplementary Fig.1-14 for "CLCNet: a contrastive learning and chromosome-aware network for genomic prediction in plants"

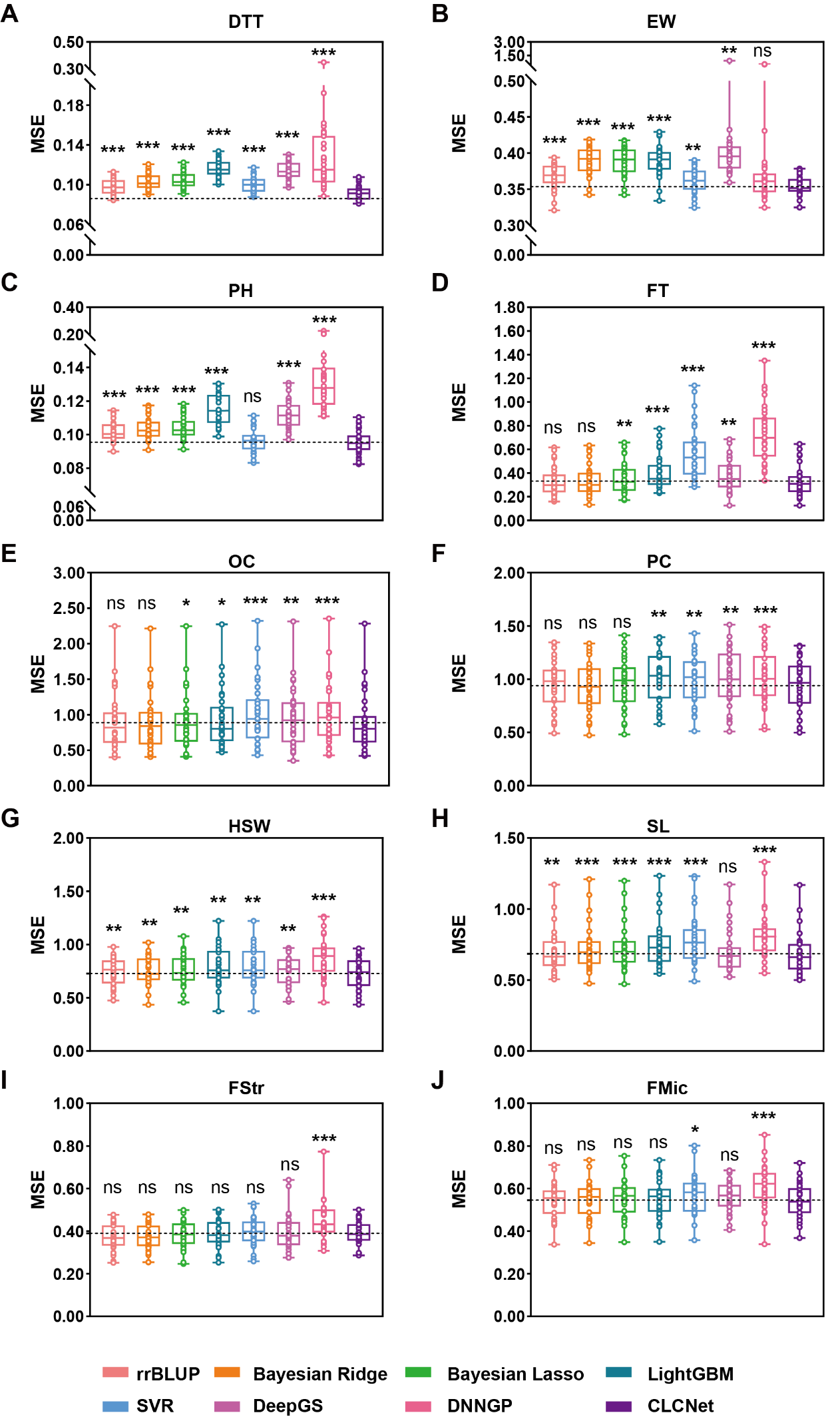

**Supplementary Fig. 1 Prediction performance of CLCNet on ten traits by MSE.**

**(A) maize DTT, (B) maize EW, (C) maize PH, (D) rapeseed FT, (E) rapeseed OC, (F) rapeseed PC, (G) soybean HSW, (H) soybean SL, (I) cotton FStr and (J) cotton FMic.** Trait abbreviations correspond to: DTT-days to tasseling, EW-ear weight, PH- plant height, FT-flowering time, OC-oil content, PC-protein content, HSW-hundred seed weight, SL-seed length, FStr-fiber strength, and FMic-fiber micronaire. MSE denoted mean squared error. The dashed line indicated the average PCC of CLCNet. A two-tailed paired t-tests with Benjamini–Hochberg false discovery rate (FDR) correction was used to compare CLCNet with other models. *** denoted *p* < 0.001, ** denoted *p* < 0.01, * denoted *p* < 0.05, and ns indicated no significant difference.

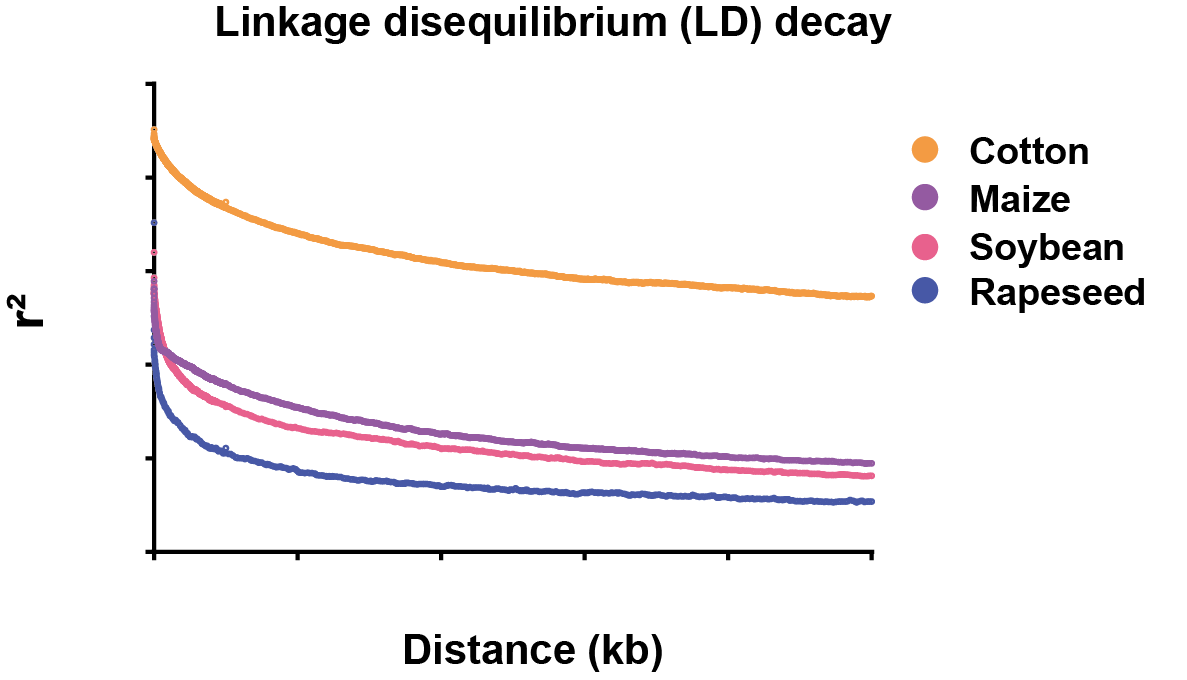

**Supplementary Fig. 2 Comparison of genome-wide linkage disequilibrium (LD) decay across species.**

**Decay of LD, quantified by the squared correlation coefficient (*r*^2^), relative to physical distance (kb) in cotton, maize, soybean, and rapeseed.**

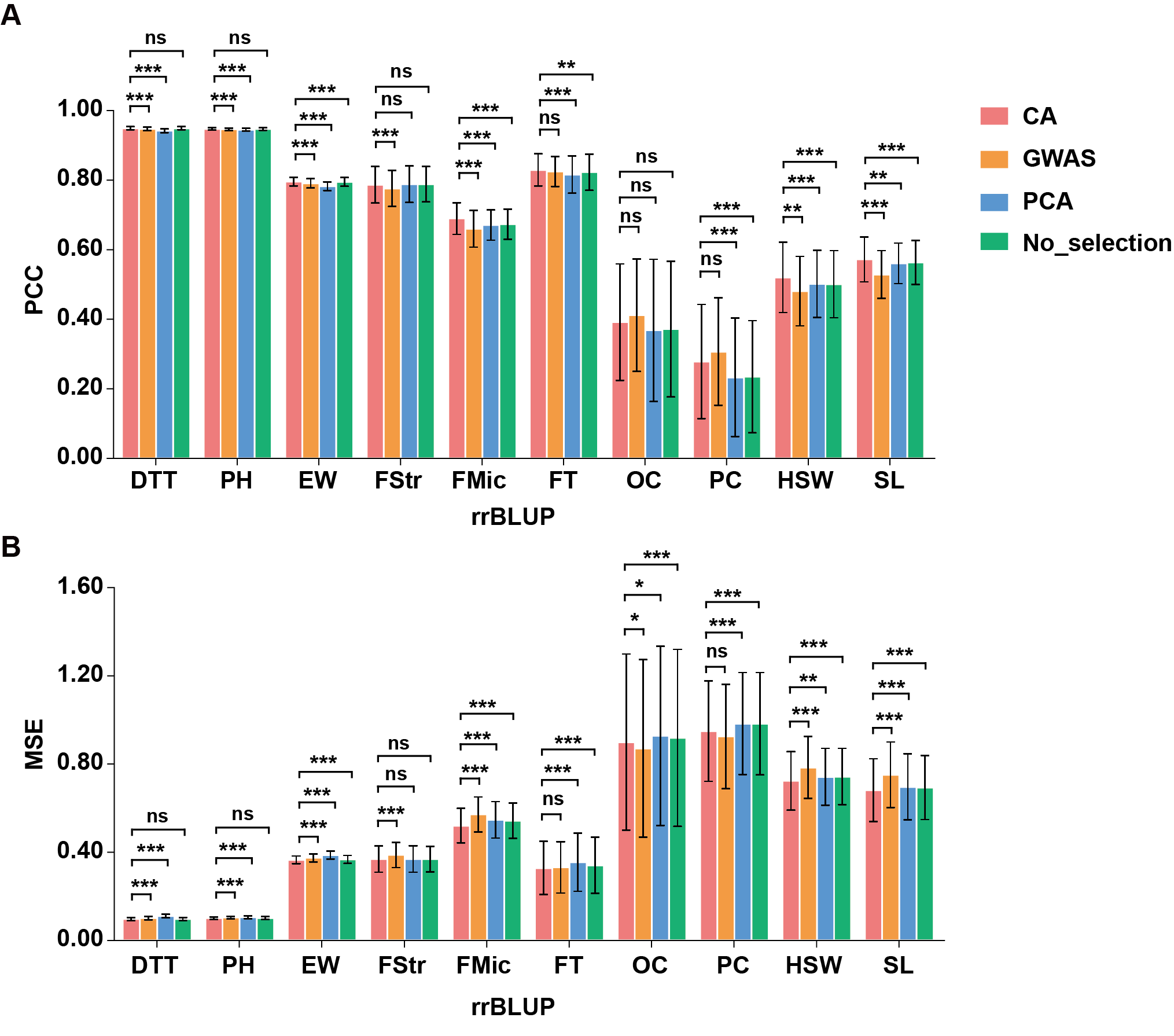

**Supplementary Fig. 3 Performance comparison of feature selection methods based on PCC and MSE in rrBLUP.**

**(A) Comparison the PCC of CA, GWAS and PCA feature selection methods. (B) Comparison the MSE of CA, GWAS and PCA feature selection methods.** A two-tailed paired t-tests with Benjamini–Hochberg false discovery rate (FDR) correction was used to compare CA with other methods. *** denoted *p* < 0.001, ** denoted *p* < 0.01, * denoted *p* < 0.05, and ns indicated no significant difference.

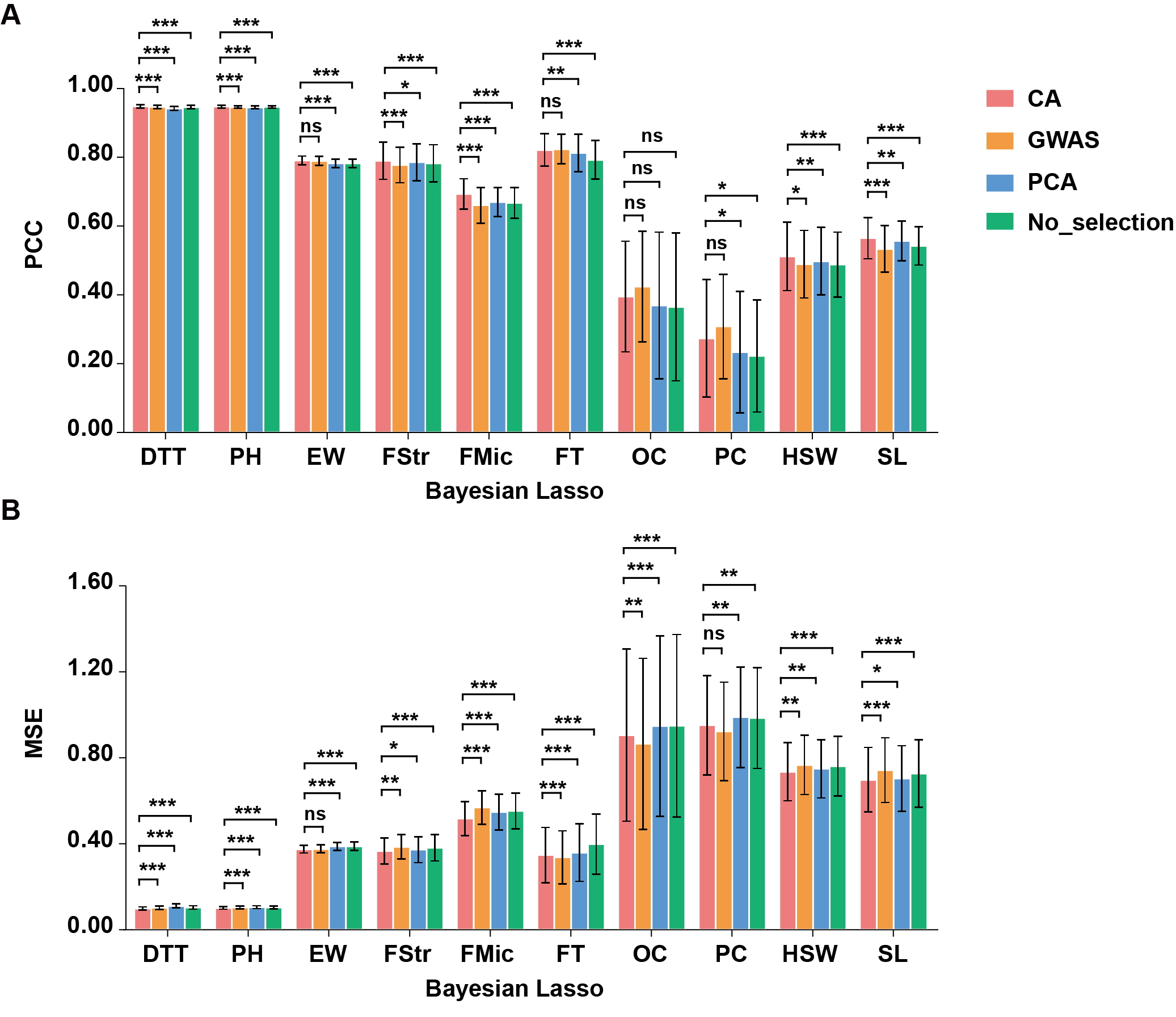

**Supplementary Fig. 4 Performance comparison of feature selection methods based on PCC and MSE in Bayesian Lasso.**

**(A) Comparison the PCC of CA, GWAS and PCA feature selection methods. (B) Comparison the MSE of CA, GWAS and PCA feature selection methods.** A two-tailed paired t-tests with Benjamini–Hochberg false discovery rate (FDR) correction was used to compare CA with other methods. *** denoted *p* < 0.001, ** denoted *p* < 0.01, * denoted *p* < 0.05, and ns indicated no significant difference.

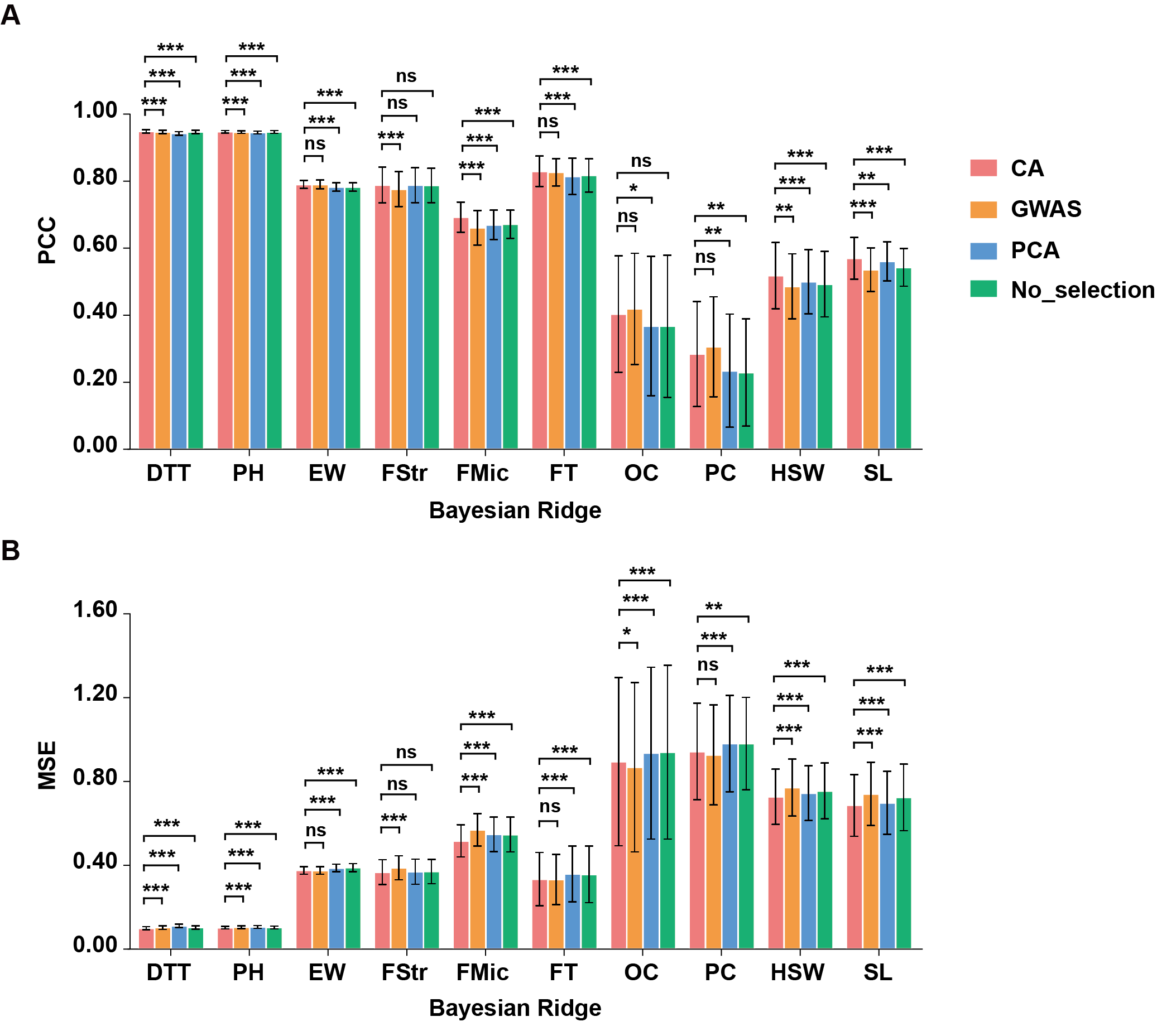

**Supplementary Fig. 5 Performance comparison of feature selection methods based on PCC and MSE in Bayesian Ridge.**

**(A) Comparison the PCC of CA, GWAS and PCA feature selection methods. (B) Comparison the MSE of CA, GWAS and PCA feature selection methods.** A two-tailed paired t-tests with Benjamini–Hochberg false discovery rate (FDR) correction was used to compare CA with other methods. *** denoted *p* < 0.001, ** denoted *p* < 0.01, * denoted *p* < 0.05, and ns indicated no significant difference.

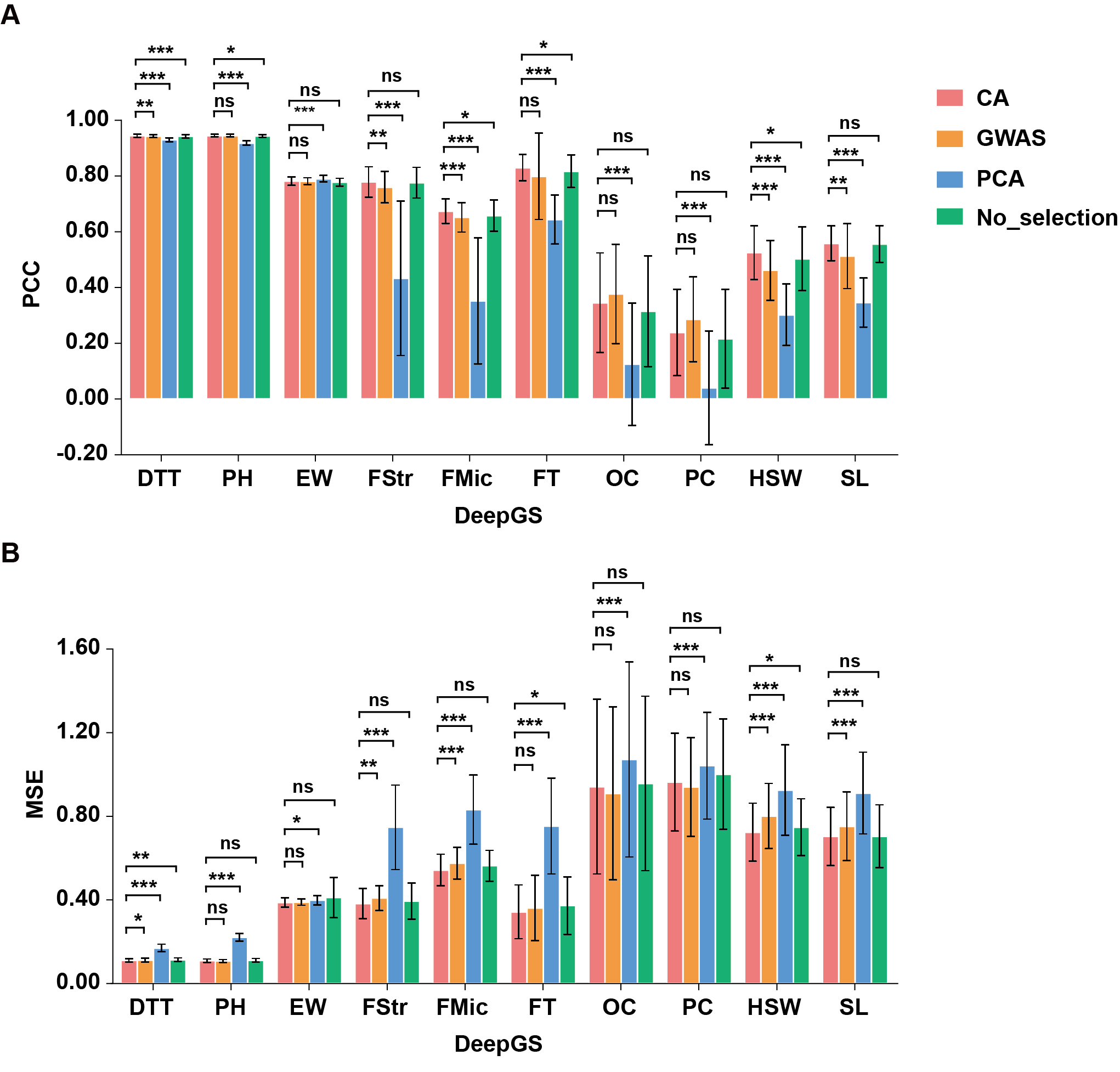

**Supplementary Fig. 6 Performance comparison of feature selection methods based on PCC and MSE in** **DeepGS.**

**(A) Comparison the PCC of CA, GWAS and PCA feature selection methods. (B) Comparison the MSE of CA, GWAS and PCA feature selection methods.** A two-tailed paired t-tests with Benjamini–Hochberg false discovery rate (FDR) correction was used to compare CA with other methods. *** denoted *p* < 0.001, ** denoted *p* < 0.01, * denoted *p* < 0.05, and ns indicated no significant difference.

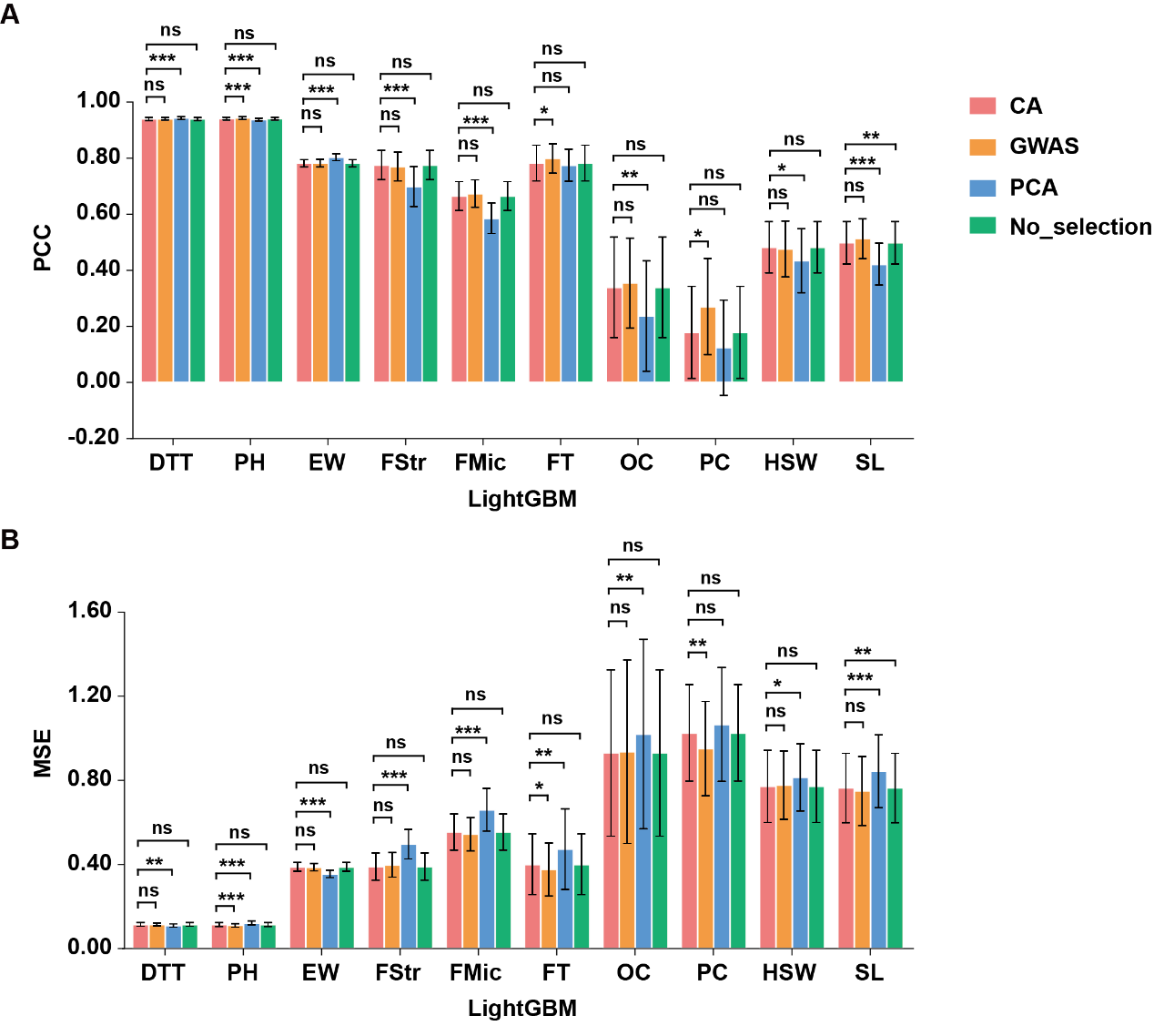

**Supplementary Fig. 7 Performance comparison of feature selection methods based on PCC and MSE in** **LightGBM.**

**(A) Comparison the PCC of CA, GWAS and PCA feature selection methods. (B) Comparison the MSE of CA, GWAS and PCA feature selection methods.** A two-tailed paired t-tests with Benjamini–Hochberg false discovery rate (FDR) correction was used to compare CA with other methods. *** denoted *p* < 0.001, ** denoted *p* < 0.01, * denoted *p* < 0.05, and ns indicated no significant difference.

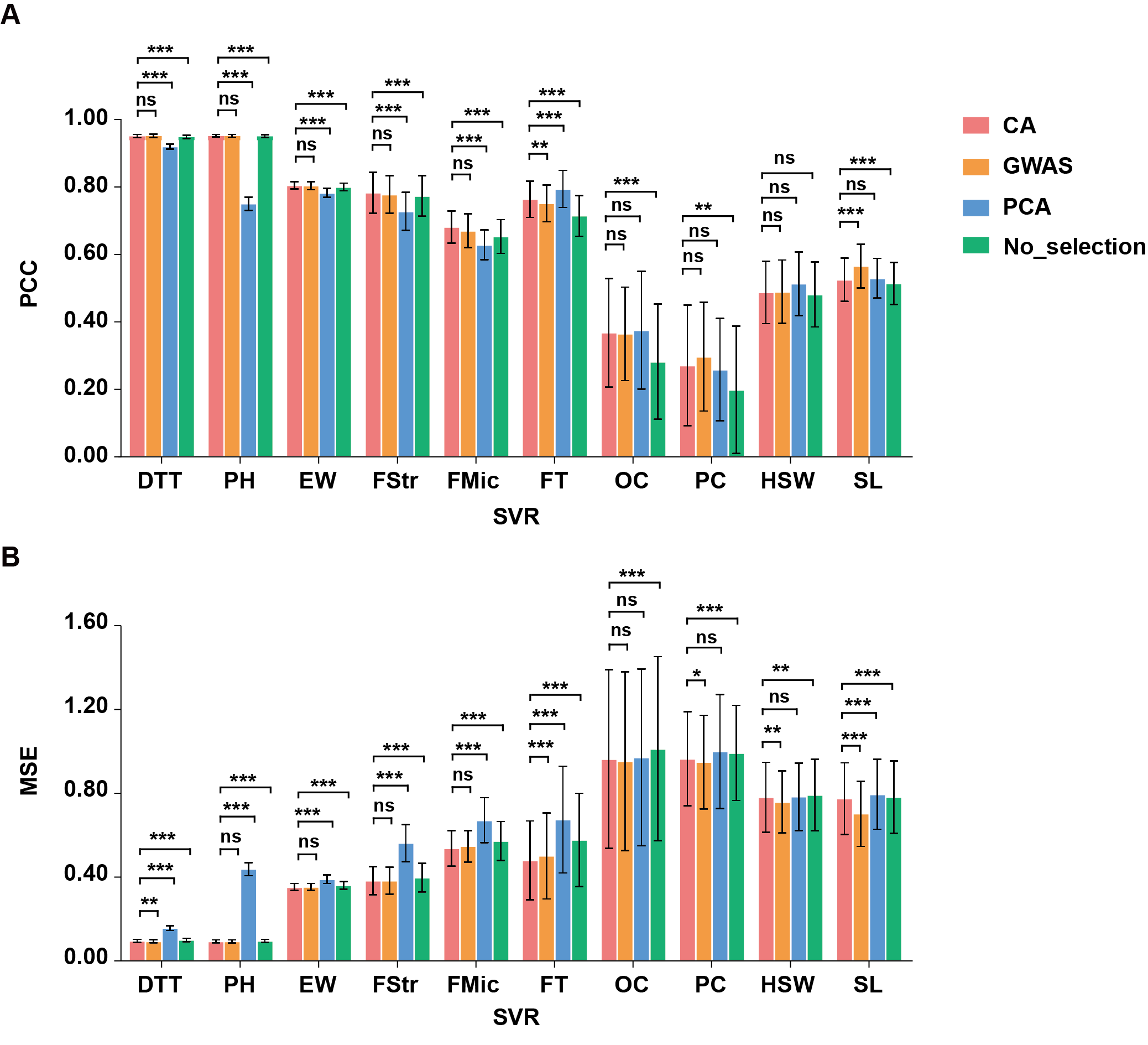

**Supplementary Fig. 8 Performance comparison of feature selection methods based on PCC and MSE in** **SVR.**

**(A) Comparison the PCC of CA, GWAS and PCA feature selection methods. (B) Comparison the MSE of CA, GWAS and PCA feature selection methods.** A two-tailed paired t-tests with Benjamini–Hochberg false discovery rate (FDR) correction was used to compare CA with other methods. *** denoted *p* < 0.001, ** denoted *p* < 0.01, * denoted *p* < 0.05, and ns indicated no significant difference.

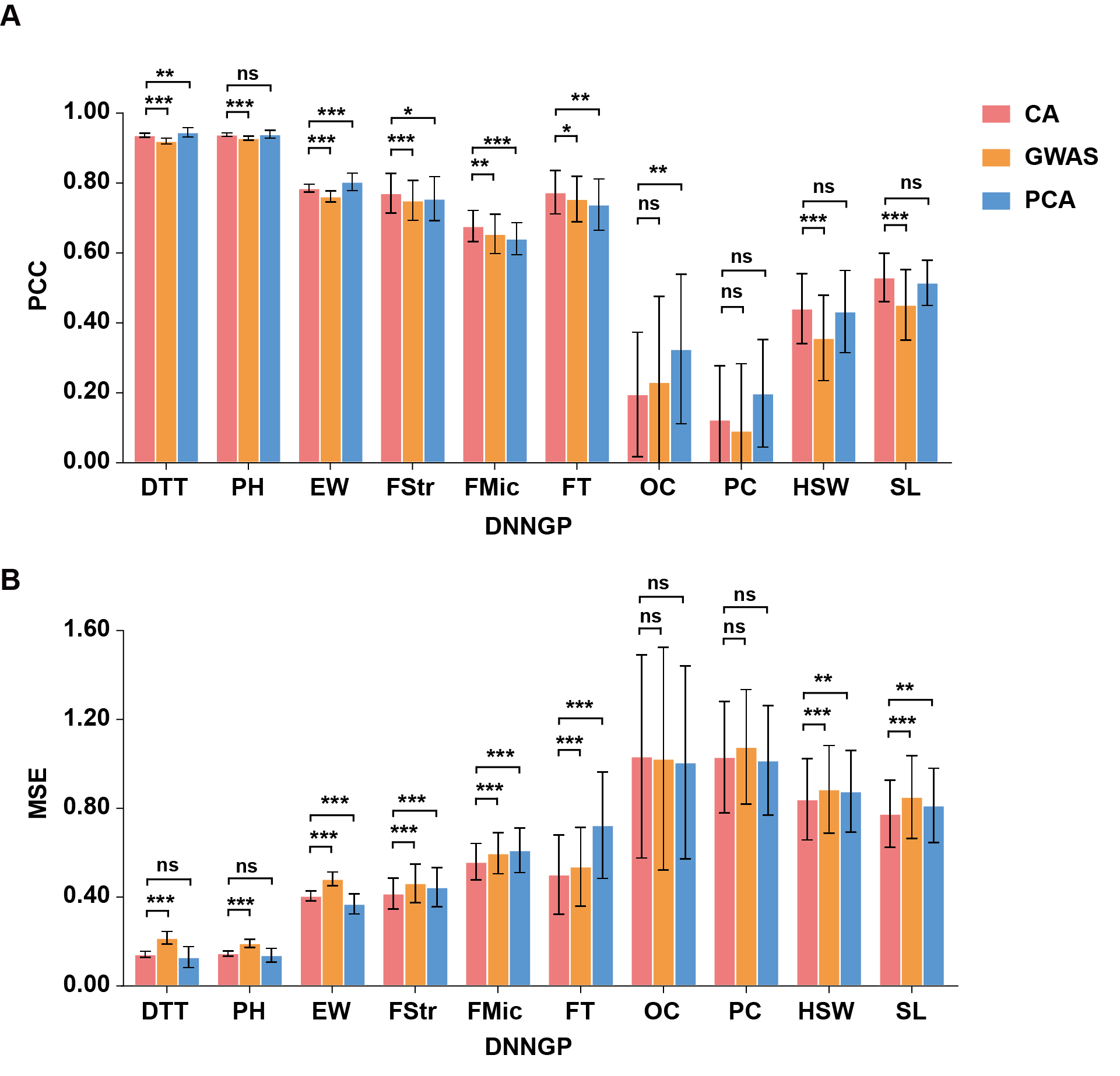

**Supplementary Fig. 9** **Performance comparison of feature selection methods based on PCC and MSE in DNNGP.**

**(A) Comparison the PCC of CA, GWAS and PCA feature selection methods. (B) Comparison the MSE of CA, GWAS and PCA feature selection methods.** A two-tailed paired t-tests with Benjamini–Hochberg false discovery rate (FDR) correction was used to compare CA with other methods. *** denoted *p* < 0.001, ** denoted *p* < 0.01, * denoted *p* < 0.05, and ns indicated no significant difference.

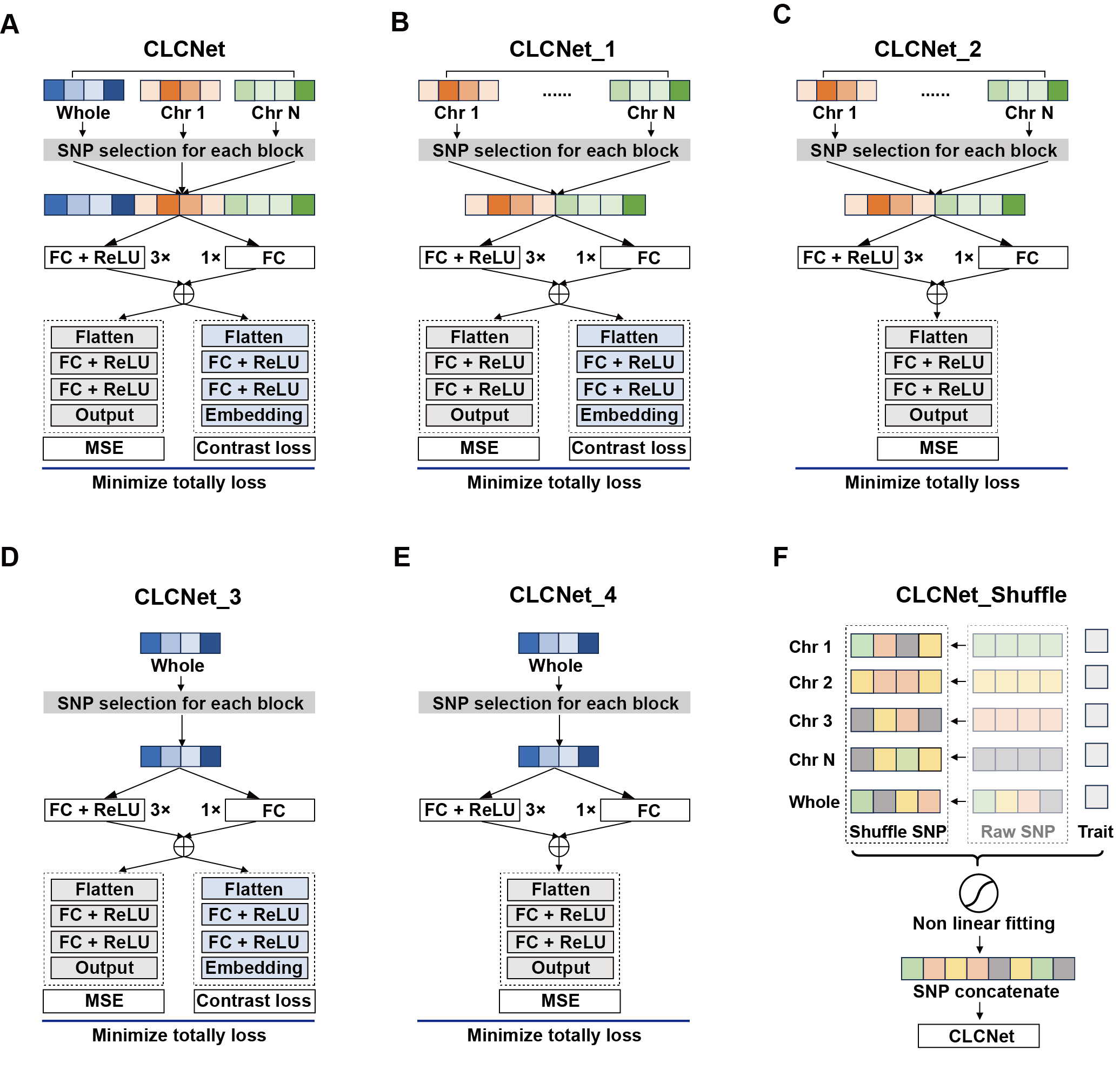

**Supplementary Fig. 10** **Model architecture diagrams.**

**(A) CLCNet model, (B) CLCNet_1, (C) CLCNet_2, (D) CLCNet_3, (E) CLCNet_4, and (E) CLCNet_Shuffle.**

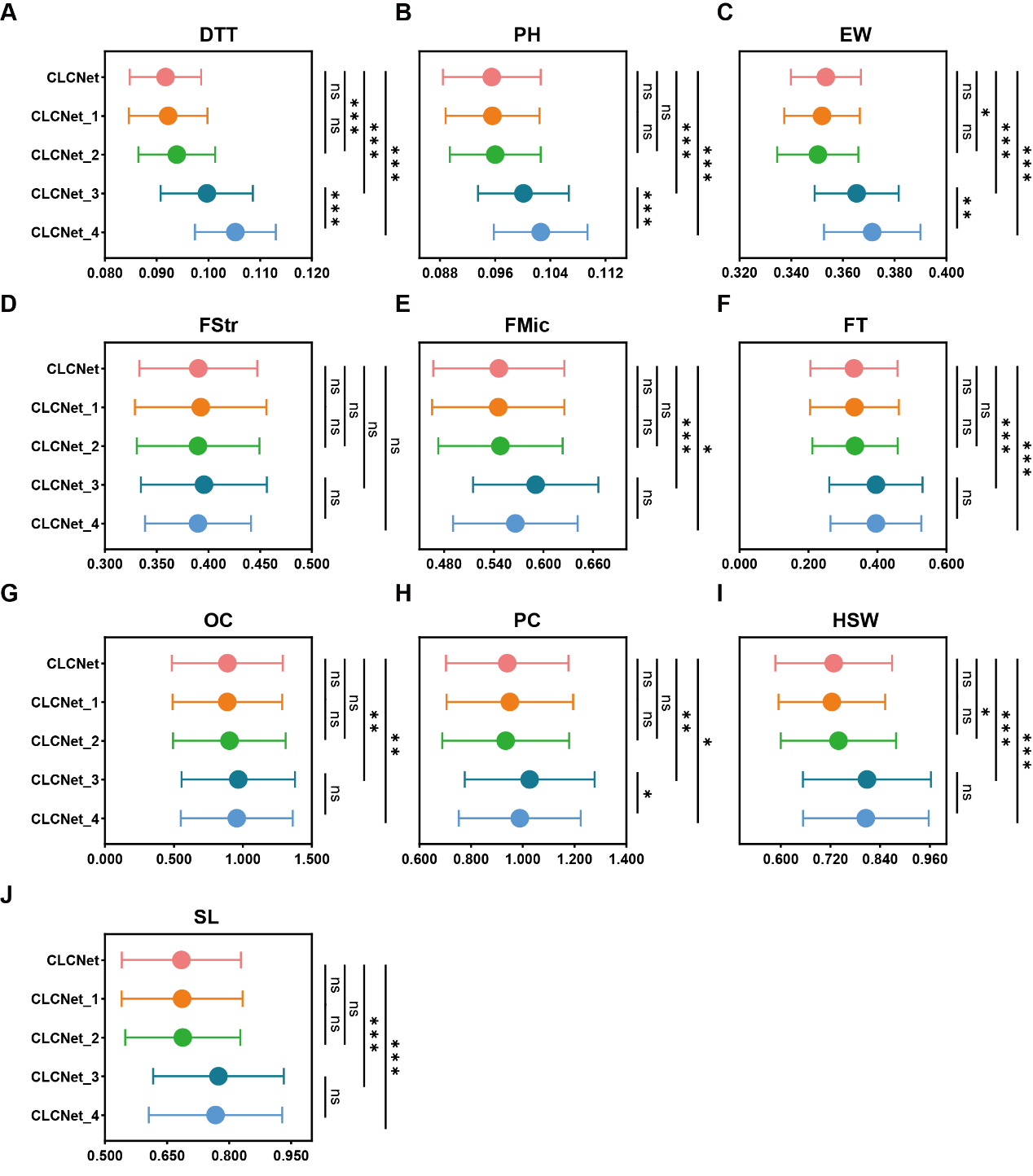

**Supplementary Fig. 11** **Performance of CLCNet's core modules on MSE.**

**(A) maize DTT, (B) maize PH, (C) maize EW, (D) cotton FStr, (E) cotton FMic, (F) rapeseed FT, (G) rapeseed OC, (H) rapeseed PC, (I) soybean HSW and (J) soybean SL.** A two-tailed paired t-tests with Benjamini–Hochberg false discovery rate (FDR) correction was used. *** denoted *p* < 0.001, ** denoted *p* < 0.01, * denoted *p* < 0.05, and ns indicated no significant difference.

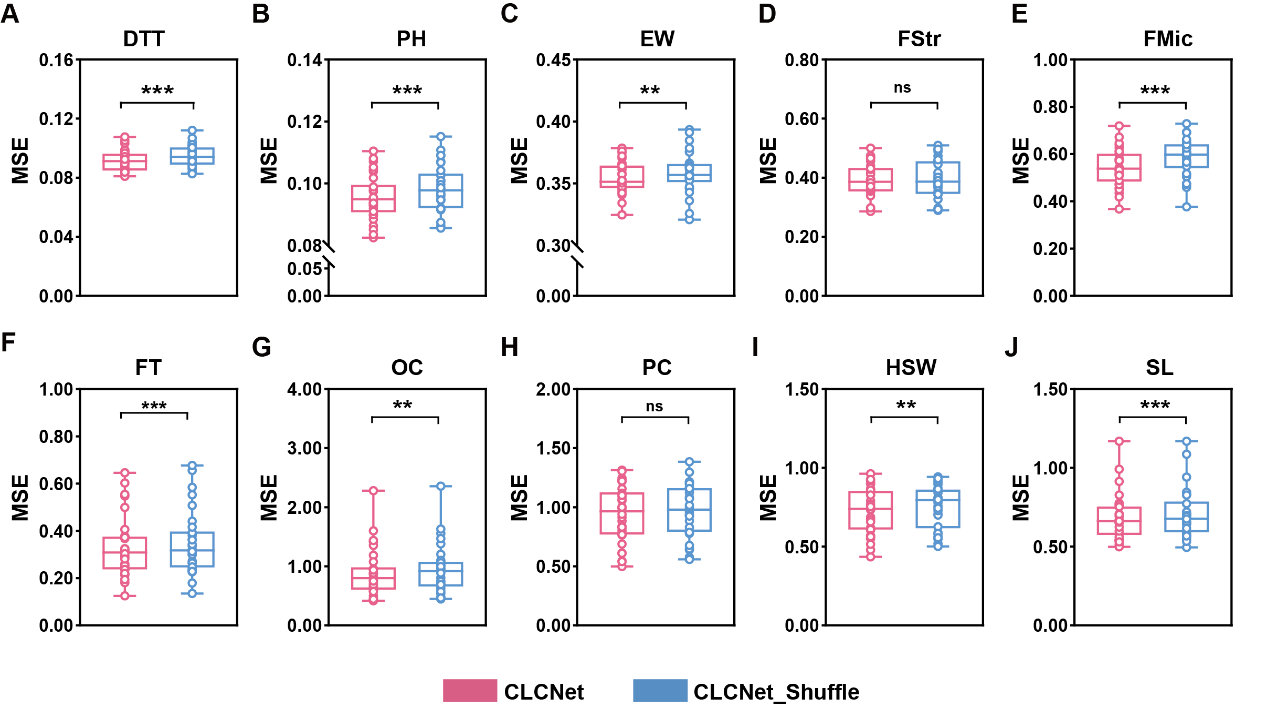

**Supplementary Fig. 12 Model MSE performance under SNP position perturbation.**

**(A) maize DTT, (B) maize PH, (C) maize EW, (D) cotton FStr, (E) cotton FMic, (F) rapeseed FT, (G) rapeseed OC, (H) rapeseed PC, (I) soybean HSW and (J) soybean SL.** A two-tailed paired t-tests with Benjamini–Hochberg false discovery rate (FDR) correction was used to CLCNet with CLCNet_Shuffle version. *** denoted *p* < 0.001, ** denoted *p* < 0.01, * denoted *p* < 0.05, and ns indicated no significant difference.

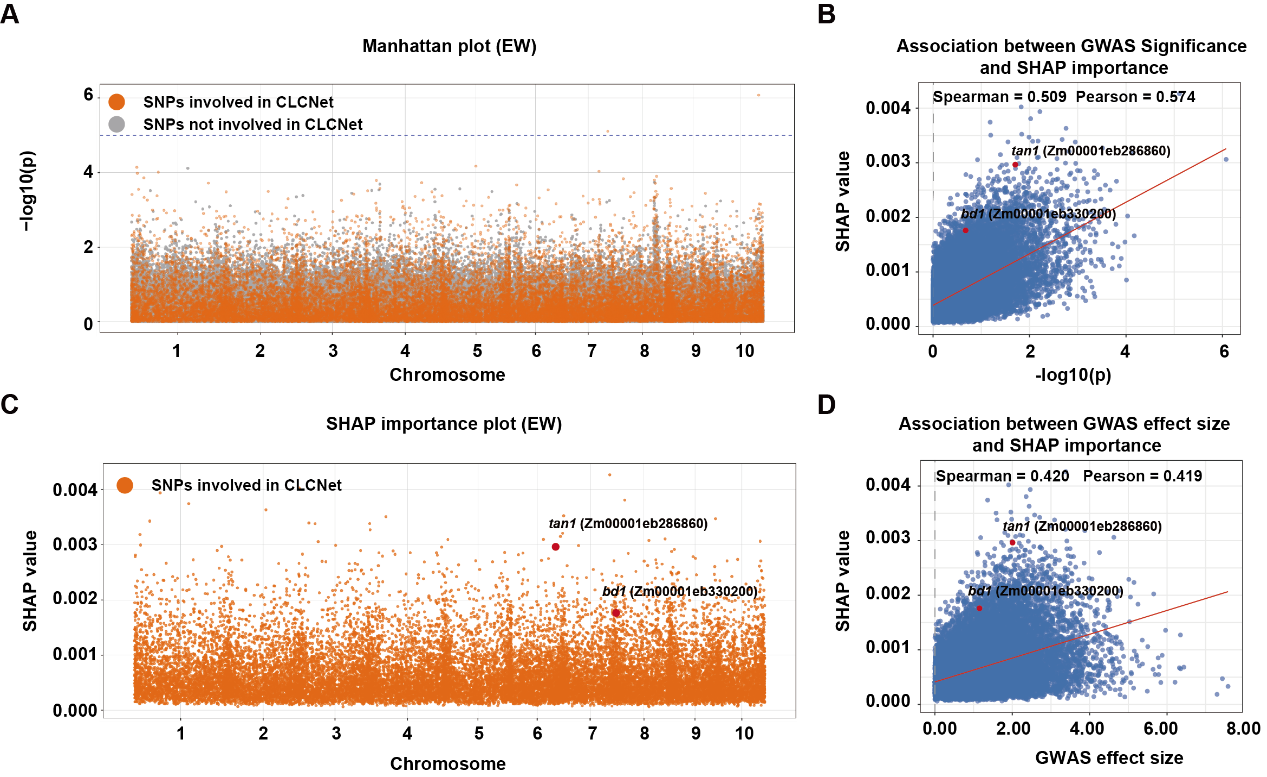

**Supplementary Fig. 13 Relationship between SHAP-based SNP importance derived from CLCNet and GWAS for EW.**

**(A) Manhattan plot of GWAS results for EW. (B) Relationship between GWAS association significance and SNP SHAP importance learned by CLCNet for EW. (C) SHAP-based SNP importance distribution derived from the CLCNet for EW. (D) Relationship between GWAS estimated effect size and SNP SHAP importance learned by CLCNet for EW.**

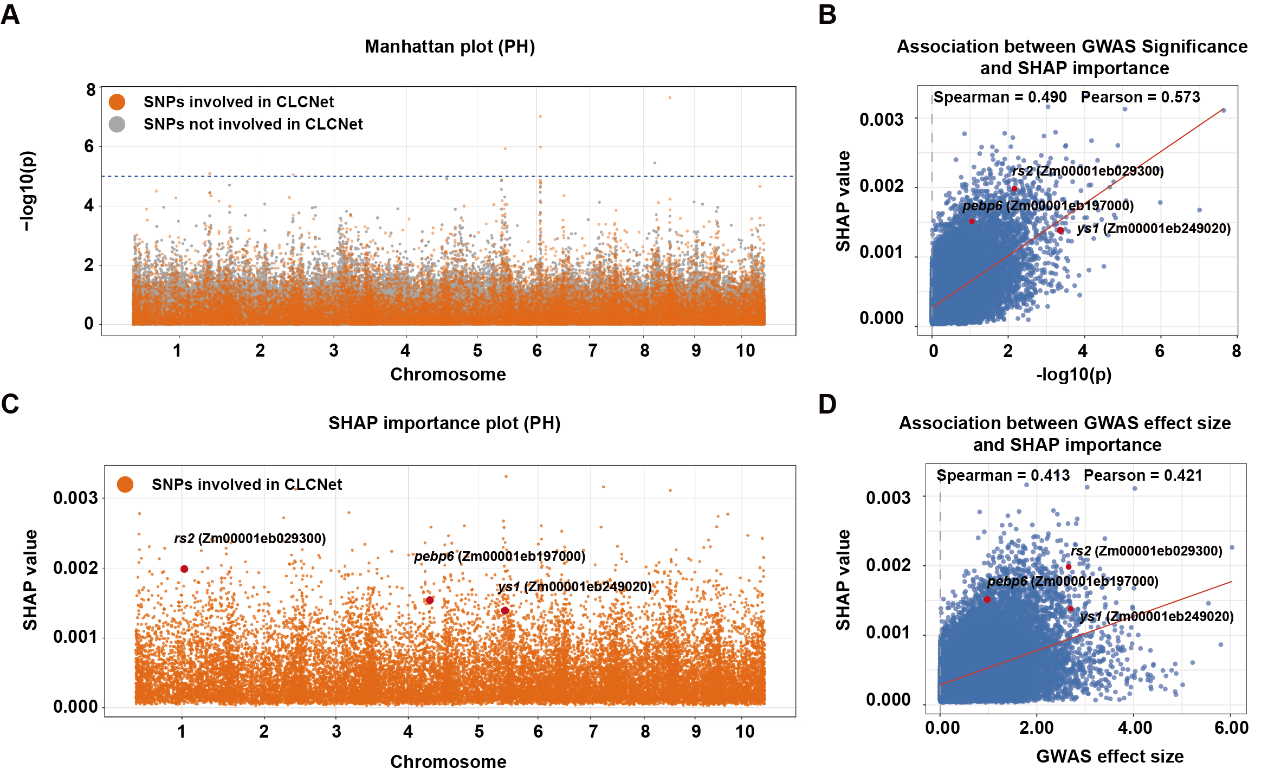

**Supplementary Fig. 14 Relationship between SHAP-based SNP importance derived from CLCNet and GWAS for PH.**

**(A) Manhattan plot of GWAS results for PH. (B) Relationship between GWAS association significance and SNP SHAP importance learned by CLCNet for PH. (C) SHAP-based SNP importance distribution derived from the CLCNet for PH. (D) Relationship between GWAS estimated effect size and SNP SHAP importance learned by CLCNet for PH.**
